# Hypermyelination Improves Strength and Detection of Neuronal Activity in the CA1 Hippocampus and Facilitates Neuroprotection in *Fus*^OL^cKO Mice

**DOI:** 10.1101/2025.06.30.662453

**Authors:** Steven M. Wellman, Kelly Guzman, Naofumi Suematsu, Teresa Thai, Te-Hsuan Tung, Camila Garcia Padilla, Sadhana Sridhar, Keying Chen, Franca Cambi, Takashi D.Y. Kozai

**Affiliations:** Department of Bioengineering, University of Pittsburgh, Pittsburgh, PA, USA, 15213; Center for Neural Basis of Cognition, Pittsburgh, PA, USA, 15213; Veterans Administration Pittsburgh, Pittsburgh, PA, USA, 15213; Department of Neurology, University of Pittsburgh, Pittsburgh, PA, USA, 15213; Center for Neuroscience, University of Pittsburgh, Pittsburgh, PA, USA, 15213; McGowan Institute of Regenerative Medicine, University of Pittsburgh, Pittsburgh, PA, USA, 15213; NeuroTech Center, University of Pittsburgh Brain Institute, Pittsburgh, PA, USA, 15213; Department of Molecular Cell Physiology, Kyoto Prefectural University of Medicine, Kyoto, Japan, 602-8566

**Author notes:** Corresponding author: Takashi D.Y. Kozai.

## Abstract

1.0.

Loss of oligodendrocytes (OLs) and myelin impairs cortical neuronal firing and network stability, whereas enhancement of oligodendrogenesis improves electrophysiological stability in cortex and, to a lesser extent, hippocampus. OLs exhibit regional heterogeneity, especially in their ability to synthesize cholesterol, a critical driver of myelin wrapping and ensheathment of axons. Conditional depletion of the Fused in sarcoma (*Fus)* gene in OLs, referred to as *Fus*^OL^cKO, increases cholesterol biosynthesis, myelin thickness, and tissue cholesterol content. We examine whether this hypermyelination alters extracellular recordings across the layers of visual cortex and the underlying hippocampal CA1 over 16 weeks. In *Fus*^OL^cKO mice, visually-evoked single-unit detectability and firing rate in CA1 increased relative to wild-type littermates, whereas cortical recordings showed no improvement. At the population level, *Fus*^OL^cKO cortex exhibited reduced firing rates and lower functional connectivity, indicating altered network dynamics. Post-mortem histology revealed higher neuron density in recorded cortex and greater excitatory synapse density in CA1 of *Fus*^OL^cKO mice suggesting region-specific neuroprotection and synaptic strengthening. These results demonstrate that cholesterol-driven hypermyelination enhances chronic hippocampal recordings while disrupting cortical network communication. Our study highlights myelin’s region-dependent roles in supporting single-cell reliability, tuning population dynamics, and maintaining circuit integrity under chronic perturbation.

**SIGNIFICANCE STATEMENT:** Myelin critically regulates neural circuit function via conduction and metabolic support. Here, we show that cholesterol-driven hypermyelination in *Fus*^OL^cKO mice augments single-unit detection and firing in hippocampal CA1 but reduces population firing and interlaminar connectivity within the cortex. These findings reveal a dual role for myelin: it can both safeguard specific circuit activity and perturb large-scale cortical communication. Understanding these dynamics is essential for designing myelin-targeted therapies in neurodegenerative disorders.

## 3.0 INTRODUCTION

Oligodendrocytes (OLs) contribute to neural circuit function by facilitating saltatory conduction and providing metabolic and neurotrophic support to axons (Bradl & Lassmann, 2010; Edgar et al., 2004; Funfschilling et al., 2012; Lappe-Siefke et al., 2003; Nave, 2010a; Philips & Rothstein, 2017). Specifically, myelin sheaths produced by OLs support the rapid transmission of neuronal activity by electrically insulating axons (Nave, 2010a; Simons & Nave, 2015). These multilamellar membranes, enriched in cholesterol and other lipids, reduce ion leakage, conserve neuronal energy and lower the energetic cost of repolarization, while directly supplying metabolic substrates such as lactate (Castelfranco & Hartline, 2015; Philips & Rothstein, 2017; Rosko, Smith, Yamazaki, & Huang, 2019). Given their role in maintaining axonal integrity, OLs and myelin are highly vulnerable to traumatic brain injury and neuroinflammation, which can induce demyelination, OL degeneration, and impaired maturation of OL precursor cells (Nave, 2010b; Pang, Cai, & Rhodes, 2003; Wellman et al., 2020; Wellman, Li, Yaxiaer, McNamara, & Kozai, 2019). Such disruptions compromise axonal health and neurodegeneration (Cheng et al., 2024; Schäffner et al., 2023), network dysfunction (Poggi et al., 2016; Wellman et al., 2020), and lead to behavioral deficits (Serra-de-Oliveira et al., 2015). Preserving or enhancing OL and myelin function may be essential for preventing axonal degeneration, maintaining network integrity, and promoting functional recovery in neurodegenerative conditions (Simons & Nave, 2015; Wellman, Cambi, & Kozai, 2018; Wellman & Kozai, 2017).

Disruption of OL function or demyelination in response to injury or inflammation negatively impacts neuronal activity and cortical signal propagation (Duncan, Simkins, & Emery, 2021; French, Reid, Mamontov, Simmons, & Grinspan, 2009; López-Muguruza & Matute, 2023). For example, cuprizone-induced demyelination degrades single-unit detectability, alters oscillatory patterns, and disrupts laminar signaling in the cortex (Wellman et al., 2020). Conversely, enhancing oligodendrocyte function, such as through pharmacological treatment of clemastine, significantly improves electrophysiological stability by promoting neuronal survival and increasing cortical connectivity (K. Chen, Cambi, & Kozai, 2023b). These findings support a critical role for OLs and myelin in sustaining long-term neuronal activity and circuit-level function, particularly in disease contexts (Wellman & Kozai, 2017).

Despite the well-established roles of OLs and myelin in regulating neuronal activity as well as their susceptibility to neuroinflammation and traumatic brain injury, the heterogeneity of oligodendrocytes across brain regions and the distinct molecular pathways by which they support neuronal electrophysiology remains poorly understood (Chamling et al., 2021; Hayashi & Suzuki, 2019; Wellman et al., 2018). Mice with conditional depletion of the Fused in sarcoma (*Fus)* gene within oligodendrocytes, termed *Fus*^OL^cKO, exhibit increased myelin thickness driven by enhanced cholesterol biosynthesis without changes in OL numbers (Guzman et al., 2020). Selectively manipulating the *Fus* gene in this way would enable better understanding of the molecular mechanisms underlying myelin regulation and their impact on electrophysiological properties of neurons across different brain regions. Furthermore, this model would be suitable to examine the effects of myelin function in isolation, without confounding effects from changes in OL density. Interestingly, different brain anatomical and functional regions show an enrichment of specific subclusters of OLs identified by single cell transcriptomics with notable differences between CA1 and cortex (Marques et al., 2016). Mice with conditional depletion of *Fus* gene within OLs exhibit enhanced novelty-induced motor function and increased exploration behavior compared to wild-type mice (Guzman et al., 2020), suggesting a functional link between hypermyelination and neuronal activity.

In this study, we used the *Fus*^OL^cKO mouse model to investigate how oligodendrocyte-specific FUS deletion affects myelin integrity in response to microelectrode implantation and its impact on sensory-evoked neuronal electrophysiology and functional connectivity within cortical and hippocampal circuits which are enriched with different subclusters of mature OLs. Electrophysiological recordings were acquired from *Fus*^OL^cKO mice and their wild-type (WT) littermate controls over a 16-week recording period. *Fus*^OL^cKO mice enhanced the detection, signal strength, and firing rates of individually recorded neurons within the CA1 region of the hippocampus, but not cortex. *Fus*^OL^cKO mice also exhibited reduced population-level firing rates, accompanied by decreased functional connectivity within the cortex, indicating a disruption in network-wide communication. Post-mortem immunohistochemistry revealed enhanced preservation of neurons within the recorded cortical area of *Fus*^OL^cKO mice compared to WT controls, suggesting that neuroprotection afforded by hypermyelination contributed to improved chronic electrophysiological recording outcomes.

## 4.0 METHODS

The use of animals in this study followed the guidelines of Institutional Animal Care and Use Committees (IACUC) of the Veterans Administration Pittsburgh Health Sciences (Animal Welfare Assurance number A3376-01) and of the University of Pittsburgh (Animal Welfare Assurance number A3187-01) and was approved by IACUC under protocols, #0321 and #15043715. All mice were bred and housed in a barrier facility with a 12-hour light/dark cycle. Food and water were available ad libitum. The *Fus*^OL^cKO line used for these studies was previously published (Guzman et al., 2020) and was maintained by breeding *Fus*^fl/fl^/Cnp^cre/+^ (referred to as *Fus*^OL^cKO) with the *Fus*^fl/fl^ littermates (referred to as WT), used as controls for the experiments. Genotyping to detect *Fus*^fl/fl^ and Cnp^cre^ was performed, as described (Guzman et al., 2020). Both males and females were used for all the analyses.

### 4.1. Surgical electrode implantation

Single-shank Michigan-style electrodes (A16-3mm-100-703-CM16LP) were inserted into the left primary monocular visual cortex (V1) of 2 month old *Fus*^OL^cKO mice (*n* = 6) and WT(*n* = 6). The probes extend from the pial surface through the V1 to the CA1 region of the hippocampus and record both spontaneous and evoked neural activity. The position in the CA1 subfield of the hippocampus was experimentally validated by visualization of the tract after acute implantation in n = 2 mice. Procedures for surgical electrode implantation were performed as described previously (T. D. Kozai, Li, et al., 2014). Briefly, an anesthetic cocktail of xylazine (7 mg/kg) and ketamine (75 mg/kg) were injected intraperitoneally (I.P.) for surgery sedation prior to fixing the animal onto a stereotaxic frame. Hair, skin, and connective tissue were removed from the top of the skull to reveal the site of surgical implantation. Vetbond adhesive was used to dry the surface of the skull prior to drilling and provide a supportive grip for a dental cement head cap. Three bone screw holes were drilled (two over both motor cortices and one over the contralateral visual cortex) prior to insertion of 4 mm long, 0.86 mm diameter stainless steel bone screws (Fine Science Tools, British Columbia, Canada) for wrapping of ground and reference wires. The ground wire was wrapped around the bone screw over the ipsilateral motor cortex while the reference wire was wrapped over the bone screws located above the contralateral motor and visual cortex. A drill-sized craniotomy positioned at 1 mm anterior to Lambda and 1.5 mm lateral to midline was made using a high-speed dental drill and 0.7 mm drill bit. Saline was periodically applied to prevent thermal damage due to drilling. A microelectrode was perpendicularly inserted at a speed of 15 mm/s using a DC motor-controller (C-863, Physik Instructmente, Karlsruhe, Germany). To record from the hippocampus, the electrode was inserted to a cortical depth of 1600 μm, visually confirming that the last contact site disappeared beneath the pial surface. A silicone elastomer (Kwik-Sil) was used to fill the craniotomy around the electrode prior to sealing with a dental cement head cap. Body temperature was maintained and monitored using a rectangular heating pad (Deltaphase isothermal pad, Braintree Scientific Inc. Braintree, MA). Mice were given an I.P. injection of ketofen (5 mg/kg) on the day of surgery and up to two days after for post-operative recovery.

### 4.2. Electrophysiological recordings

Electrophysiological recordings were conducted inside a grounded Faraday cage to prevent electrical interference from environmental noise as described previously (Alba, Du, Catt, Kozai, & Cui, 2015; Kolarcik et al., 2015; T. D. Kozai et al., 2015; Takashi D Y Kozai et al., 2016; Nicolai et al., 2018). Mice were situated on a rotating platform for awake, head-fixed recording. Trials to obtain spontaneous neural activity were recorded in a dark room. Visually-evoked neural activity was stimulated using the MATLAB-based Psychophysics toolbox on a 24” LCD (V243H, Acer. Xizhi, New Taipei City, Taiwan) monitor located 20 cm from the contralateral eye of the mouse covering a 60° wide by 60° high visual field. A drifting gradient of solid white and black bars were presented and synchronized with the neural recording system (RX7, Tucker-Davis Technologies, Alachua, FL) at a sample rate of 24,414 Hz. Each white and black grating was presented for 1 s (rotated in 135° increments), separated by 1 s of a dark screen, and repeated for a total of 64 trials per recording session. Total recording was over 16 weeks, with daily activity measured for the first 14 days, followed by weekly recording for the remaining 14 weeks.

### 4.3. Electrophysiological data processing

#### 4.3.1. Current source density

Current source density (CSD) was used to identify layer 4 along the length of the electrode following evoked activity within the visual cortex. CSD plots were generated by computing the average evoked (stimulus-locked) LFP for each electrode site, smoothing the signal across all electrode sites, and then calculating the second spatial derivative. CSDs were averaged across 64 stimulus trials and layer 4 was identified as an inversion in LFP polarity within the first 100 ms following stimulus onset. Cortical drift and the magnitude change in drift of implanted arrays over time was reported as the average change in layer 4 depth relative to depth at day 0 (day of surgery). All electrophysiological evaluation between different cortical depths occurred following alignment of all animals in each group for each day to their corresponding layer 4 depth. Hippocampal recording was identified by the depth and the presence of activity below the electrically silent white matter tracts.

#### 4.3.2. Single-unit (SU) sorting and analysis

Processing of raw signal data occurred offline using a custom MATLAB script modified from previously published methods (T. D. Kozai et al., 2015). Data was passed through a Butterworth filter with a passband from 2 to 300 Hz to produce data containing information on local field potentials (LFPs) and 0.3 to 5 kHz to isolate spiking information. Common average referencing was applied to the data as previously described (Ludwig et al., 2009). A fixed threshold value of 3.5 standard deviations below the mean was used to identify potential neuronal single-unit (SU) and multi-unit (MU) activity. Only channels which exhibited a signal-to-noise ratio (SNR) >2 were considered for single-unit sorting. Signal-to-noise ratios (SNR) were calculated by dividing the peak-to-peak amplitude of each single unit by the noise and reported as average SNR and average SNR per active site (electrode channels reporting detection of SU) over time. Sortable single units were confirmed by observing the quality and shape of neuronal waveforms, auto-correlograms, and peri-stimulus time histograms (PSTH) with 50 ms bins. SU yield was calculated as the percentage of electrode sites (out of 16) with at least one identifiable single unit. The noise floor for each electrode site was taken as two times the standard deviation (2*STD) of the entire recorded data stream after removing all threshold crossing events. Finally, the SNR of any channels without a sortable SU waveform were reported as zero for the purposes of calculating averages.

Putative excitatory and inhibitory units were classified based on the trough-to-peak interval (TPI) as previously described (K. Chen, Forrest, Gonzalez Burgos, & Kozai, 2024). According to our whole dataset with the Gaussian mixture model fitting, we decided to use a threshold of 0.51 ms to separate units into excitatory (TPI > 0.51 ms, n = 2934) and inhibitory ones (TPI < 0.51 ms, n = 853). After the excitatory and inhibitory units were classified, unit yield (ratio of recording sites identifying excitatory/inhibitory units in total channel counts) was calculated for each animal and each day.

Signal-to-noise firing rate ratio (SNFRR) for each unit was calculated as described in previous publications (T. D. Y. Kozai, Z. Du, et al., 2015):

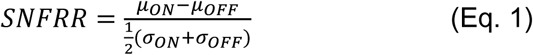

where *μ*_*ON*_ and *μ*_*OFF*_ indicate average firing rate and standard deviation (across 64 trials) during ON (stimulation) and OFF (pre-stimulation) periods. SNFRR is a normalized firing rate change relative to each spontaneous firing rate, and its positive value corresponds to an increase in firing rate by visual stimulation. To evaluate the modulation magnitude of neural activity by visual stimulation, |SNFRR| was used.

#### 4.3.3. Multi-unit (MU) analysis

Multi-unit activity was measured as any threshold crossing event that occurred within the 1 s period after each stimulus-locked trigger or pseudotrigger was recorded. Peri-stimulus time histograms (PSTHs) of 50-ms bin size were generated to gauge the dynamics of multi-unit activity. Multi-unit firing rate was measured as the average number of threshold crossing events within a 1-s period after each stimulus presentation or pseudotrigger. Evoked multi-unit activity was evaluated by calculating multi-unit yield and SNFRR. Parameters for multi-unit spike counts involved varying the temporal bin size and latency after stimulus presentation from 0 to 1 seconds in length via 1-ms increments in order to evaluate multi-unit yield and SNFRR. Multi-unit yield was defined as the number of electrode sites that had a significantly different spike count (*p* < 0.05 determined by a paired t-test) for a given bin size and latency following stimulus presentation (stim ON) compared to spike counts within that same bin size immediately before stimulation (stim OFF). SNFRR measured the difference in the firing rate of multi-unit activity before and after stimulus relative to the average standard deviation between each stimulus condition (Eq. 1).

#### 4.3.4. Laminar coherence analysis

Electrophysiological activity within and between different laminar depths was evaluated by calculating the magnitude-squared coherence, described previously (Michelson & Kozai, 2018). Coherence is a quantitative description of the similarity between two signals based on their frequency-dependent responses. Coherence was reported as a value between 0 and 1, with 0 indicating no relationship and 1 indicating a perfect linear relationship between two corresponding signals. Coherence was calculated as follows:

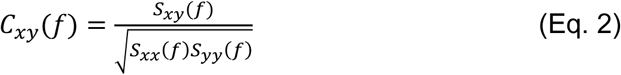

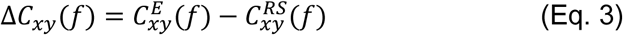

where *C*_*xy*_(*f*) is the coherence, *S*_*xy*_(*f*) is the cross spectrum, and *S*_*xx*_(*f*) and *S*_*yy*_(*f*) are the autospectra of LFP activity between electrode sites *x* and *y*, respectively. Normalized coherence Δ*C*_*xy*_(*f*) was calculated by taking the difference between evoked coherence 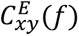 and resting-state coherence 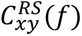. Coherence calculations were performed within 1 s intervals following each stimulus presentation or pseudotrigger, at a half-bandwidth of 3 Hz and a taper number of 5, and then averaged across all trials. Coherence between different cortical regions was assessed by comparing the average coherence of all electrode sites located within each respective layer. Coherence values for a specific frequency band were reported as the average coherence within that frequency range across all animals within each group.

#### 4.3.5. Phase-amplitude coupling (PAC) analysis

To quantify the directionality of neuronal circuit connections, phase amplitude coupling (PAC) was calculated based on Kullback-Leibler (KL) formula as previously described (Koch et al., 2020; Tort, Komorowski, Eichenbaum, & Kopell, 2010). The raw signal was processed with a bandpass filtered for specific local field potential frequencies *f*. The Hilbert transform was applied to output the time series of the phase component from the slow frequency LFP activity in Channel X, denoted as *ϕ*_*X*_(*t*, *f*_*X*_) where *t* represents time and *f*_*X*_ represents phase frequencies 2,3,4… 20Hz. The time series of the amplitude component at high frequency LFP activity in channel Y ( *A*_F_(*t*, *f*_F_) ) was extracted using Morlet wavelet transform (5 cycles) per frequency, using amplitude frequencies *f*_F_ = 30,31,32…90Hz.

Next, we calculated the amplitude of high frequency LFP oscillation in Channel Y at each given slow frequency LFP activity in Channel X, resulting in the composite time series *ϕ*_*X*_(*t*, *f*_*X*_), *A*_*Y*_(*t*, *f*_*Y*_). The phases were binned every 18° (N = total number of phase bins, 20) and the amplitude at each phase bin (*i*) was averaged to give *A*_*Y*_(*t*, *f*_*Y*_)_*ϕ*X(*t*,*f*_^X^_)_(*i*). Lastly, normalizing by the sum of all bins, resulted in the normalized amplitude distribution over phases *P*(*i*, *f*_*X*_, *f*_*Y*_).

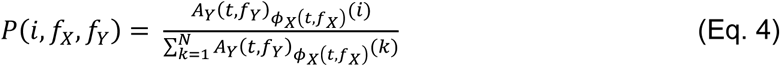

If there is no PAC between channels X and Y, the normalized amplitude distribution *P*(*i*, *f*_*X*_, *f*_*Y*_) would result in a uniform distribution. Further, the joint entropy defined as *H*(*f*_*X*_, *f*_*Y*_) measures the amount of deviation of the uniform distribution given by the following:

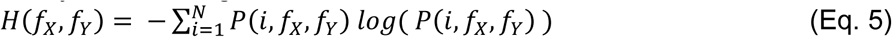

At the maximum joint entropy *H*_*o*_ = *logN* ^2^, the normalized amplitude distribution *P*(*i*, *f*_*X*_, *f*_*Y*_) is uniform. We then calculated the difference between *H*(*f*_*X*_, *f*_*Y*_) and *H*_*o*_ using the KL distance formula. The modulation index (MI) of PAC was then calculated by dividing the KL distance by the uniform distribution *H*_*o*_.

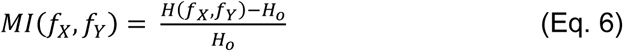

When Channel X = Channel Y, a higher MI value of PAC indicates stronger LFP synchronization coupling between the low frequency phase and high frequency amplitude and increased activation level of the intralaminar network. When Channel X ≠ Channel Y, a larger MI value of PAC suggests enhanced LFP synchronization coupling between the low frequency phase in the upstream layer and the high frequency amplitude in the downstream layer, resulting in an increase in the directional interlaminar circuitry.

### 4.4. Electrochemical impedance spectroscopy

Electrochemical impedances were measured prior to each recording session. Mice were awake and head-fixed on a rotating platform while the implanted electrode was connected to an Autolab potentiostat along with a 16-channel multiplexer. Impedances were measured from each channel using a 10 mV RMS sine wave ranging from 10 Hz to 32 kHz. Impedances were reported as the average impedance at 1 kHz over all animals for each day unless otherwise noted.

### 4.5. Immunohistochemistry

Mouse brains were intracardially perfused with 4% paraformaldehyde (PFA), post-fixed in 4% PFA for 18 hours, and kept in 30% sucrose at 4°C until samples achieved equilibration by sinking to the bottom of the solution. Brains were frozen while embedding with optimum cutting temperature (OCT) media directly on the headpiece of the cryostat (Leica) and sectioned horizontally at 14 μm thickness starting from the convexity of the brain down to 800 μm below the cortex. To evaluate the CA1 subregions 14 μm sections were collected between 900 μm and 1600 μm. To help identify the probe tract n=2 per genotype were sectioned between 2000 to 2300 μm. Sections adhered to positively charged slides were permeabilized and blocked using 0.1% Triton-X with 10% normal goat serum in PBS at room temperature for 1 hr and incubated with primary antibodies to CC1 (1:100, Millipore, Burlington, Massachusetts USA, #OP80), Olig2 (1:200 Millipore, Burlington, Massachusetts USA, #Mabn50), NG2 (1:100, Millipore, Burlington, Massachusetts USA, #Ab5320), GFAP (1:1000 Abcam, Cambridge, Massachusetts USA, #Ab4674), Iba-1 (1:1000 Wako, Richmond, Virginia USA, #019–19741), MBP (1:500 Abcam, Cambridge, Massachusetts USA, #Ab40390), NeuN (1:200 Millipore, Burlington, Massachusetts USA, #MAB377), Neurofilament (NF-200) (1:200, Sigma-Aldrich, St. Louis, Missouri USA # N4142-.2ML),), vGlut1 (1:200 Sigma Aldrich, St. Louis Missouri, USA, #Ab5905) and PSD95 (1:300, Abcam, Cambridge, Massachusetts, USA, #Ab18258 ) overnight at 4°C. Alexa Fluor 488/594 conjugated secondary Ab (1:300 Jackson ImmunoResearch, West Grove, Pennsylvania USA # 115-585-003, # 111-605-003, #111-545-003), IgG2b isotope specific (1:300 anti-mouse 488 Jackson ImmunoResearch, West Grove, Pennsylvania USA #115-545-207) and 633 anti-chicken (1:300 Thermo Fisher Scientific, Waltham, Massachusetts #a21103) were reacted for 1 hr at RT. Sections were washed and allowed to dry overnight in the dark before mounting with Fluoromount-G and DAPI (SouthernBiotech, Birmingham, Alabama USA, #0100-20). For synaptic staining, to keep the refractive index uniform through the section and surrounding materials we used Pro Long Glass as mounting agent which has a refractive index of 1.552 after 48hrs of curing (ThermoFisher Scientific, Waltham, Massachusetts, USA, #P36982) and 1.5H coverslips (Thor Labs CG15KH1).

### 4.6. Imaging and data analysis

For cellular markers, 20x TIFF images were captured over the probe site and at an equivalent area in the contralateral hemisphere or baseline tissue with a Nikon TiE epifluorescent microscope (Melville, NY USA) with NIS Element software. For the horizontal sections, grey scale, individual layer images were cropped to square and loaded into using a previously published I.N.T.E.N.S.I.T.Y. MATLAB script where the binning was applied around the probe site (Du et al., 2017; T. D. Kozai, Gugel, et al., 2014). Three bins were applied 0-50 μm, 50-100 μm and 100-150 μm extending from the center of the probe site or image of non-implanted tissue. Labeled cells were counted per bin at 400-800 μm through the cortical depth per brain. For intensity analysis, images were acquired by collecting 20x z-stack fields with 9 steps (1 μm step size) using Nikon A1R Confocal microscope (Melville, NY USA) with NIS Elements software. The sum intensity projections (SUMIP) were processed through MATLAB by applying 10 μm bins up to 15 bins concentrically around the site of probe insertion, contralateral tissue, or baseline tissue in non-implanted mice. Data were expressed as the fold intensity change at the probe site over the intensity in the corresponding area in the contralateral hemisphere (T. D. Kozai, Gugel, et al., 2014).

Excitatory synapses were quantified by counting appositions of pre-synaptic puncta stained with vGlut1 and post-synaptic puncta stained with PSD95 that are <200 nm apart in the CA1 (Hong et al., 2016). This analysis was performed on SIM (Structure Illumination Microscopy, Nikon A1R) super-resolution images (Hong, Wilton, Stevens, & Richardson, 2017) using a workflow in the Nikon General Analysis (GA3) software, which was developed in collaboration with Nikon software specialist.

### 4.7. Statistics

Changes in recording metrics with respect to time were modeled using a linear mixed effects model, described previously (Eles et al., 2017). To fit the model to nonlinear relationships, a restricted cubic spline was implemented by placing 4 knots at the 5^th^, 35^th^, 65^th^, and 95^th^ percentiles of the data. Group (*Fus*^OL^cKO and WT) and group-by-time interactions were included as fixed effects. A likelihood ratio test was performed on the estimated model. Confidence intervals were determined using a bootstrapping method with 1000 iterations. A 95% confidence interval was taken as 1.96 times the standard error of the model output. For histological analysis, a one-way ANOVA or t-test (*p* < 0.05) was used to determine significant differences in tissue stains between *Fus*^OL^cKO and WT mice. Pairwise significances were determined using a post-hoc t-test followed by a Bonferroni correction, to reduce the probability of type I error when performing multiple comparisons.

## 5.0 RESULTS

To assess how hypermyelination following OL-specific deletion of the *Fus* gene affects neuronal electrophysiology and local circuit connectivity, extracellular recordings were acquired from intracortical microelectrodes implanted in *Fus*^OL^cKO mice (**Fig. 1A**). A 16-channel Michigan-style linear electrode array was implanted to a depth of 1.6 mm, targeting both the primary visual cortex (V1) and the CA1 subregion of the hippocampus. Wild-type (WT) littermates were similarly implanted and served as controls. Recordings were collected at least once a week over a 16-week recording period. Immunohistochemistry of explanted brain tissue at 2 and 16-weeks post-implantation was performed to contextualize electrophysiological data within the biological context in the electrode-tissue microenvironment. Current source density (CSD) in response to sensory stimulation was used to monitor electrode position by tracking the cortical layer 4 sink over time (**Fig. 1B**). Electrode drift was minimal after the first post-implantation week, with deviations limited to ∼100 μm, allowing consistent depth-resolved analyses of intra- and interlaminar activity throughout the recording period.

**Figure 1.**
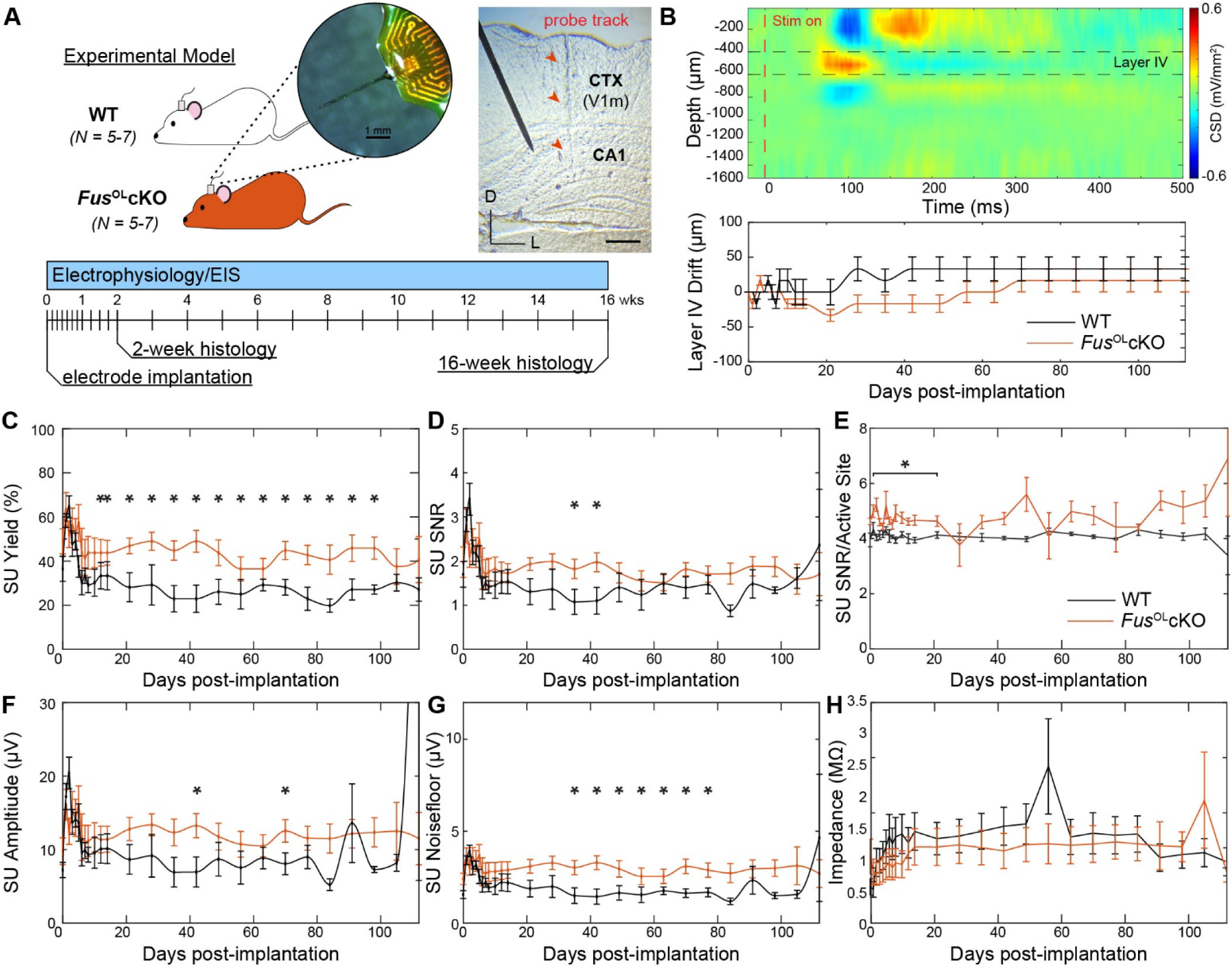
*Fus*^OL^cKO mice improve detection of individual neurons over a 16-week recording period. (A) Schematic of experimental models used and timeline of electrophysiological recording and explant histology. Inset depicts 1.6 mm long, laminar 16-channel microelectrode array. *Right*: Image of a coronal tissue section taken through a light microscope of probe track (*red arrowheads*) extending through V1 cortex and CA1 region of the hippocampus. Scale bar = 500 μm. (B) Representative current source density (CSD) plot demonstrating 500ms of electrical current activity versus electrode implant depth following visual stimulus presentation (‘stim on’) in the visual cortex used to identify layer 4 depth (sink = red, source = blue). Average drift of layer 4 depth along implanted microelectrode shank compared to day 0 of electrode insertion between WT (*black solid*) and *Fus*^OL^cKO mice (*orange solid*). (C-H) Single-unit (SU) electrophysiological metrics over time between WT (*black solid*) and *Fus*^OL^cKO mice (*orange solid*). (C) Single-unit yield over time. (D) Single-unit SNR (mean peak-peak amplitude over 2*STD of noise floor). (E) SNR per electrode channel detecting a single unit (active site). (F) Single-unit amplitude over time. (G) Amplitude of noise floor over time. (H) Average impedance reported at 1 kHz over time. * indicates non-overlapping 95% confidence intervals at each time point as determined by likelihood ratio test applied to a linear mixed effects model for WT and *Fus*^OL^cKO mice.

Both *Fus*^OL^cKO and WT mice exhibited an initial decline in electrophysiological performance, consistent with prior reports using similar intracortical electrodes. Recordings are typically most robust during the first post-implantation week, followed by signal attenuation associated with local neuron loss and gliosis, and eventual stabilization over time (K. Chen et al., 2023b; T. D. Kozai, Li, et al., 2014; Wellman et al., 2020). This biphasic trajectory reflects early optimal signal capture followed by dynamic changes at the electrode-tissue interface. When averaged across all depths, *Fus*^OL^cKO mice showed a significantly greater percentage of recording sites detecting discrete neuronal waveforms compared to WT controls between 10- and 98-d post-implantation (**Fig. 1C**, p < 0.05, likelihood ratio test with non-overlapping 95% confidence intervals). Signal-to-noise ratios (SNR) of individual waveforms were also significantly increased between 35- and 42-d post-implantation of *Fus*^OL^cKO mice compared to WT controls (**Fig. 1D**, p < 0.05, likelihood ratio test with non-overlapping 95% confidence intervals).

To more directly evaluate the quality of neuronal recordings independent of electrode failure or glial encapsulation, we next focused the analysis on ‘active sites’, defined as electrode channels that detected at least one discrete neuronal waveform on a given recording day. This subset isolates the functional interface between the electrode and viable neurons, minimizing confounds from silent channels or tissue damage. *Fus*^OL^cKO mice exhibited significantly higher SNR values at active sites compared to WT controls (**Fig. 1E**, p < 0.05, likelihood ratio test with non-overlapping 95% confidence intervals). The improved SNR was primarily driven by elevated signal amplitudes, which were significantly increased in *Fus*^OL^cKO mice at 42- and 70-d post-implantation (**Fig. 1F**, p < 0.05, likelihood ratio test with non-overlapping 95% confidence intervals). Additionally, amplitude of the noise floor was significantly elevated in *Fus*^OL^cKO mice between 35- and 77-d post-implantation (**Fig. 1G**, p < 0.05, likelihood ratio test with non-overlapping 95% confidence intervals). However, electrochemical impedance spectroscopy revealed no significant differences in impedance magnitude across the 16-week period between *Fus*^OL^cKO and WT controls (**Fig. 1H**, p < 0.05, likelihood ratio test with non-overlapping 95% confidence intervals), indicating that electrode functionality and the degree of glial encapsulation were comparable. Because Johnson–Nyquist noise scales with impedance, and impedance was equivalent across groups, elevated noise floor in *Fus*^OL^cKO mice is unlikely to arise from increased thermal noise or encapsulation. Instead, this pattern suggests a shift in the neuronal source population contributing to the broadband signal: hypermyelination may preserve proximal neuronal viability and support increased spontaneous activity or synaptic input, leading to greater asynchronous summation from nearby neurons. These findings collectively support the conclusion that *Fus*^OL^cKO enhances the long-term fidelity of neuronal recordings by improving neuron-electrode coupling through preserved myelin integrity and circuit viability.

### 5.1. Fus^OL^cKO mice improve the strength and detection of neurons within CA1 region of the hippocampus

Aberrant myelination is a hallmark of multiple neurodegenerative disorders and contributes to network dysfunction, yet the consequences of hypermyelination on neuronal signal detection remain poorly understood. To determine how OL-specific *Fus* deletion affects neuronal firing properties across brain regions, we analyzed the same electrophysiological metrics separately in V1 and CA1 subregion of the hippocampus. Previous work demonstrated that cuprizone-induced demyelination reduces extracellular signal strength and single-unit detection compared to non-demyelinated controls (Wellman et al., 2020). Since myelination alters the efficiency of action potential propagation and may influence axonal excitability and synchrony, we hypothesized that increasing myelin thickness in *Fus*^OL^cKO mice would enhance the detectability and strength of extracellularly recorded spikes. *Fus*^OL^cKO mice exhibited a significantly higher percentage of recording sites detecting discrete neuronal waveforms in CA1, but not cortex, compared to WT controls during the first week of recording as well as at 42- and 49-d post-implantation (**Fig. 2A**, p < 0.05, likelihood ratio test with non-overlapping 95% confidence intervals). Similarly, SNR of individual waveforms in CA1 were significantly increased between 35- and 49-d post-implantation in *Fus*^OL^cKO mice relative to WT (**Fig. 2B**, p < 0.05, likelihood ratio test with non-overlapping 95% confidence intervals). This SNR increase resulted from elevated signal amplitudes in CA1 of *Fus*^OL^cKO mice, which were significantly greater at both 35- and 42-d post-implantation (**Fig. 2C**, p < 0.05, likelihood ratio test with non-overlapping 95% confidence intervals). Furthermore, noise floor amplitude was significantly increased in CA1 of *Fus*^OL^cKO mice between 35- and 77-d post-implantation compared to WT (**Fig. 2D**, p < 0.05, likelihood ratio test with non-overlapping 95% confidence intervals). No significant differences in waveform properties were observed in cortex between groups, indicating that hypermyelination in *Fus*^OL^cKO mice predominantly affects neuronal activity in hippocampal CA1.

**Figure 2.**
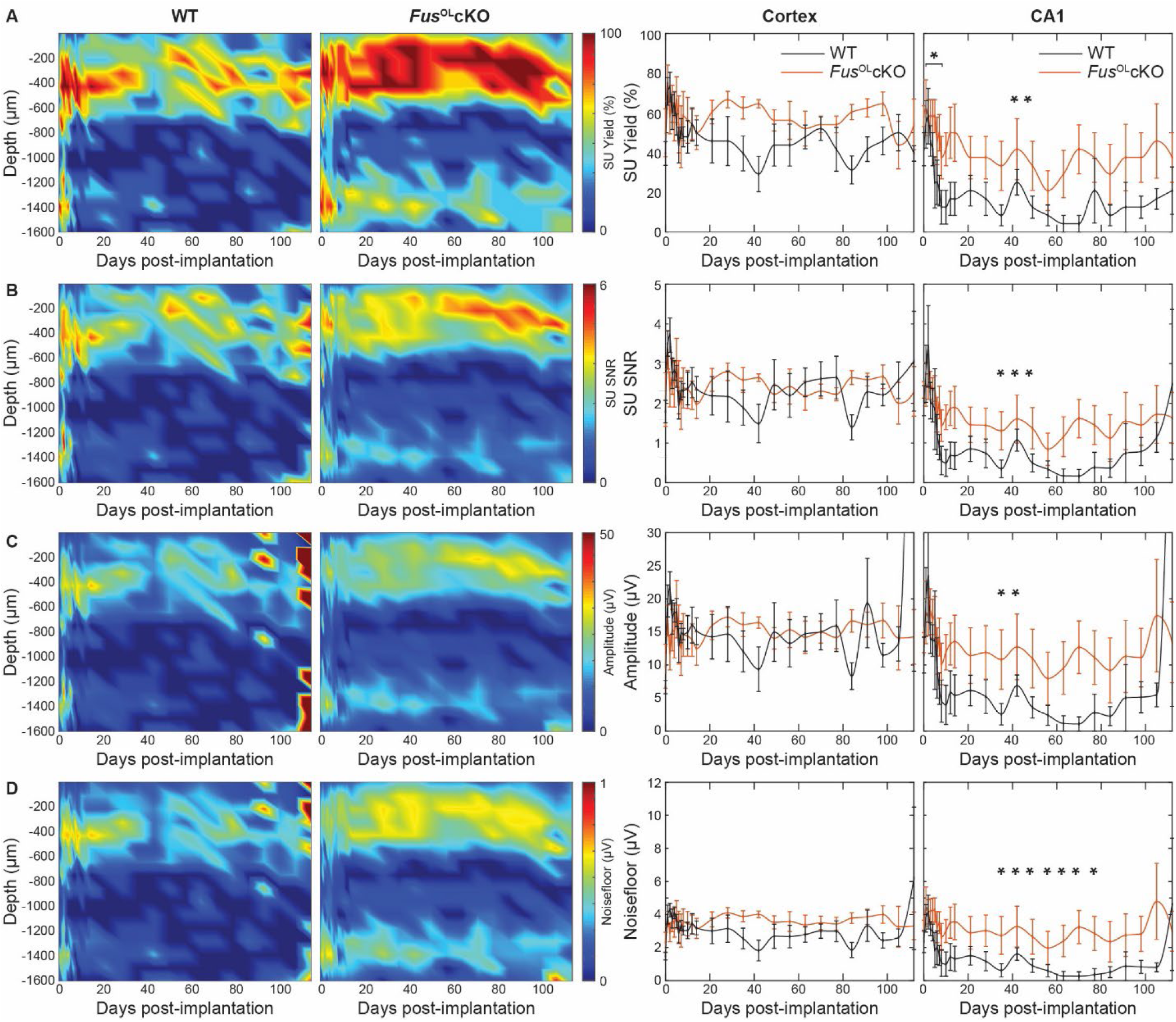
FusOLcKO mice exhibit enhanced strength and detection of individual neuronal waveforms within the CA1 region of the hippocampus. Single-unit (SU) metrics were plotted as averages over time and depth for FusOLcKO mice and WT controls. Line plots demonstrate comparable differences in single-unit quality between FusOLcKO mice and WT controls in a region-specific manner. (A) Recording yield is significantly increased in CA1 region of the hippocampus in FusOLcKO mice compared to WT controls. (B) Signal-to-noise (SNR) of single-unit activity significantly increased in CA1 region of the hippocampus in FusOLcKO mice compared to WT controls. (C) The amplitude of single-unit recordings significantly increased in CA1 region of the hippocampus in FusOLcKO mice compared to WT controls. (D) The noisefloor of single-unit recordings significantly increased in CA1 region of the hippocampus in FusOLcKO mice compared to WT controls. * indicates non-overlapping 95% confidence intervals at each time point as determined by likelihood ratio test applied to a linear mixed effects model for WT and FusOLcKO mice.

### 5.2. Fus^OL^cKO mice enhance spontaneous and evoked firing rates of individual neurons within CA1 region of the hippocampus

The degree of oligodendrocyte-mediated myelination influences not only conduction velocity but also energy efficiency and reliability of axonal transmission, especially in long-range projections such as those targeting the hippocampus. Previous studies demonstrated that cuprizone-induced demyelination reduces firing rates of individual neurons, while pharmacological preservation of oligodendrocytes and myelin maintains firing over long-term recordings (K. Chen et al., 2023b; Wellman et al., 2020). Therefore, we hypothesized that hypermyelination in *Fus*^OL^cKO mice would increase evoked neuronal firing rates compared to WT controls. Heatmaps of neuronal activity revealed depth-dependent differences in both spontaneous and visually evoked firing rates between *Fus*^OL^cKO and WT mice (**Fig. 3A,C**). Quantification of neuronal firing rates showed no significant differences between groups in cortex (**Fig. 3B**, p > 0.05, likelihood ratio test with non-overlapping 95% confidence intervals). In contrast, spontaneous firing rates in the CA1 hippocampal region were significantly elevated in Fus^OL^cKO mice compared to WT across the chronic recording period from 7- to 16-wks post-implantation (p < 0.05, likelihood ratio test with non-overlapping 95% confidence intervals). Similarly, visually evoked firing rates were unchanged in cortex but significantly increased in CA1 of *Fus*^OL^cKO mice relative to WT both acutely during the first week and chronically from 7 to 16 weeks post-implantation (**Fig. 3D**, p < 0.05, likelihood ratio test with non-overlapping 95% confidence intervals). These findings suggest that hypermyelination facilitates not only signal detection but also intrinsic and stimulus-driven excitability in hippocampal neurons, potentially reflecting improved conduction fidelity or reduced metabolic constraints. Together these results indicate that oligodendrocyte-specific *Fus* deletion enhances both spontaneous and evoked firing of hippocampal neurons over prolonged periods.

**Figure 3.**
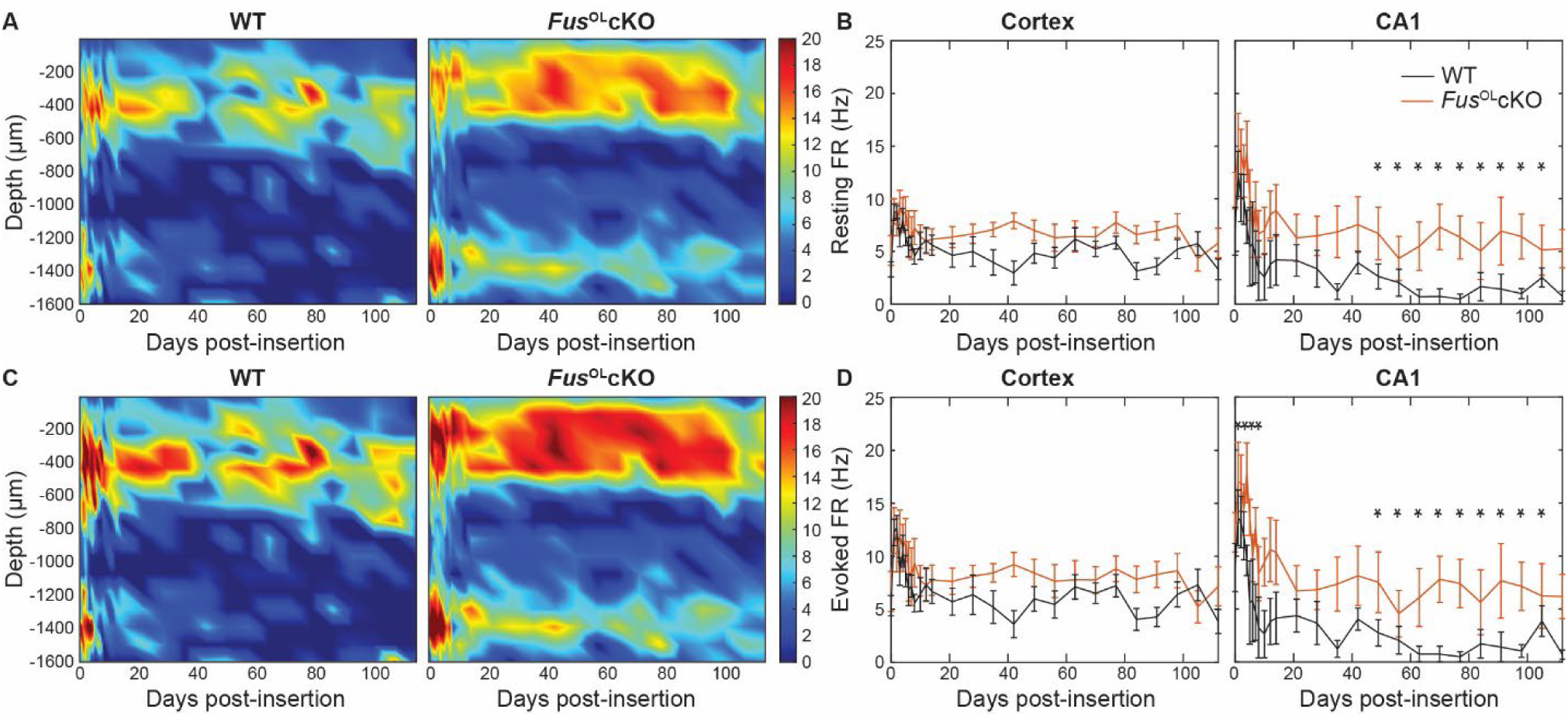
*Fus*^OL^cKO mice demonstrate elevated firing rates of individual neurons within the CA1 region of the hippocampus. Heatmaps demonstrate the average firing rate between WT and *Fus*^OL^cKO mice as a function of depth and time during (A) resting and (C) visually evoked activity. Average firing rates between WT and *Fus*^OL^cKO mice were quantified separately within cortex and subcortical CA1 region of the hippocampus for (B) resting and (D) visually evoked activity. <u>Firing rate activity significantly increased in both resting and evoked</u> states in the CA1 region of *Fus*^OL^cKO mice but not in the cortex. **\*** indicates non-overlapping 95% confidence intervals at each time point as determined by likelihood ratio test applied to a linear mixed effects model for WT and *Fus*^OL^cKO mice.

### 5.3. Fus^OL^cKO mice preserve excitatory yield and early inhibitory responsiveness in a region- and cell type-specific manner

Oligodendrocytes influence not only axonal conduction but also long-term circuit stability, particularly in excitatory-inhibitory (E/I) networks. However, how hypermyelination affects the yield and responsiveness of specific neuronal subtypes over time remains poorly defined. To investigate how hypermyelination influences neuronal subtype-specific activity, we separated putative excitatory and inhibitory units based on the trough-to-peak interval (TPI) of their spike waveforms. A Gaussian mixture model was fit to the TPI distribution, identifying a classification threshold of 0.51 ms: units with TPI > 0.51 ms were categorized as putative excitatory, while those with TPI < 0.51 ms were categorized as putative inhibitory (**Fig. 4A**, left). Representative waveforms for each category (mean ± 1 SD) are shown (**Fig. 4A**, right). Average yield of putative excitatory and inhibitory units across depth revealed a significantly higher putative excitatory unit yield in *Fus*^OL^cKO mice compared to WT controls, while putative inhibitory unit yield was greater in WT mice compared to *Fus*^OL^cKO (**Fig. 4B**, p < 0.05, likelihood ratio test with non-overlapping 95% confidence intervals). This suggests that hypermyelination may improve the detection, stability, or viability of excitatory neurons while having a more limited or suppressive effect on inhibitory unit detection.

**Figure 4.**
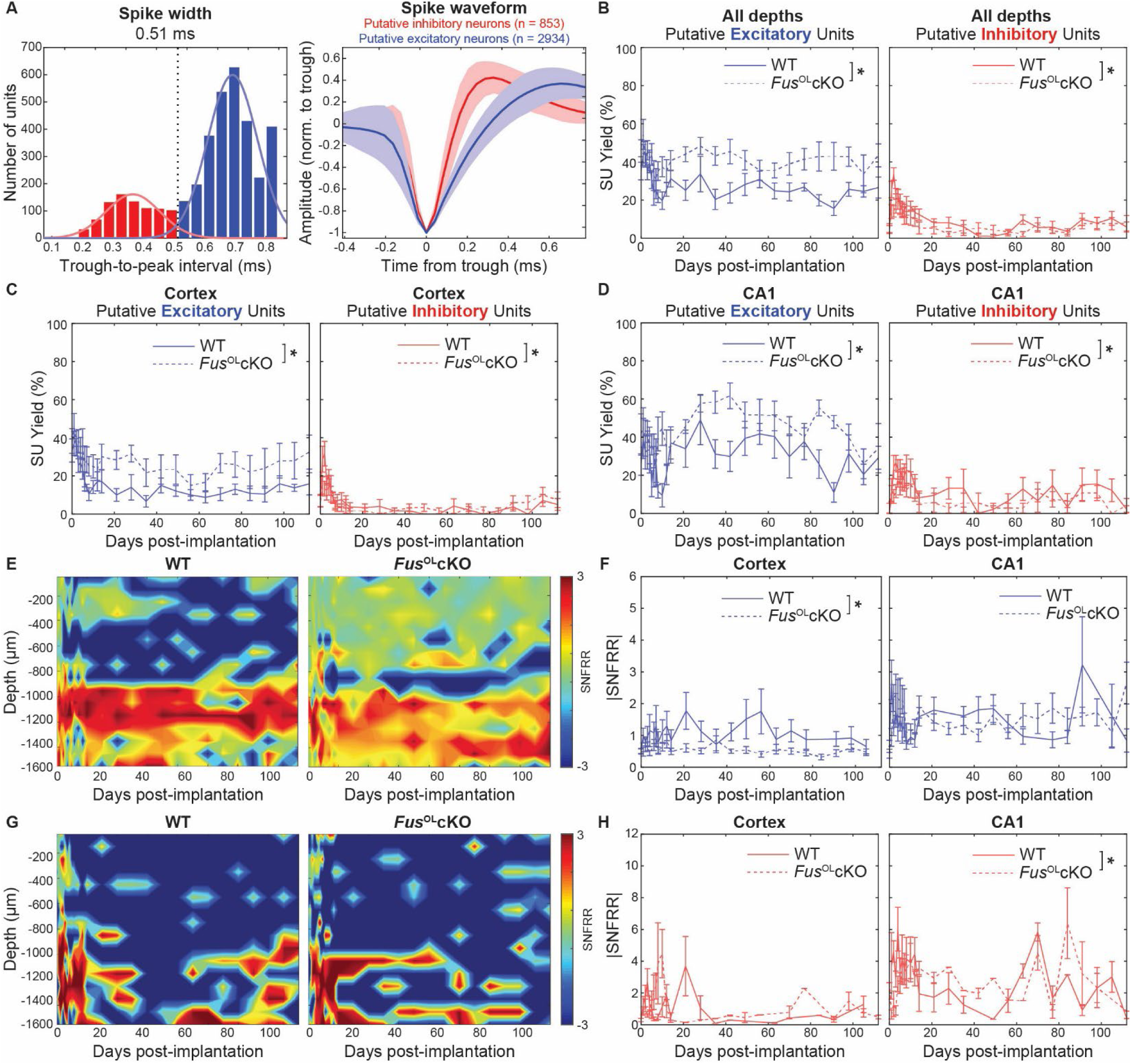
*Fus*^OL^cKO mice show selectively preserved excitatory neuron yield and responsiveness over time and across brain regions. (A) Classification of putative excitatory and inhibitory units based on trough-to-peak interval (TPI). Left: Gaussian mixture model applied to TPI distribution to determine classification threshold (0.51 ms). Right: Representative average spike waveforms (mean ± 1 SD) for excitatory (blue, TPI > 0.51 ms) and inhibitory (red, TPI < 0.51 ms) units. (B) Longitudinal analysis of excitatory and inhibitory unit yield over entire recording depth between *Fus*^OL^cKO and WT mice. (C) Longitudinal analysis of excitatory and inhibitory unit yield within the cortex between *Fus*^OL^cKO and WT mice. Excitatory yield declined over time in WT but was significantly preserved in *Fus*^OL^cKO mice. Inhibitory yield declined between both groups with limited protection conferred by hypermyelination. (D) Excitatory and inhibitory unit yield in CA1 region of hippocampus between *Fus*^OL^cKO and WT mice. Excitatory yield was preserved in *Fus*^OL^cKO mice relative to WT mice. Inhibitory yield declined over time, with WT mice showing greater yield early in the recording period. (E) Heatmap of signal-to-noise firing rate ratio (SNFRR) of excitatory units in the cortex and in CA1. (F) Average |SNFRR| of excitatory units within the cortex and CA1 region of the hippocampus in *Fus*^OL^cKO and WT mice. (G) Heatmap of signal-to-noise firing rate ratio (SNFRR) of inhibitory units in the cortex and in CA1. (H) Average |SNFRR| of inhibitory units within the cortex and CA1 region of the hippocampus in *Fus*^OL^cKO and WT mice. Inhibitory |SNFRR| in CA1 declined over time, with *Fus*^OL^cKO young mice showing transient early protection.

To determine whether these differences varied across brain regions, we analyzed longitudinal unit yield in cortex and hippocampal CA1. Putative excitatory unit yield in cortex declined over time in WT mice (**Fig. 4C**, left), consistent with glial scarring or electrode drift. In contrast, *Fus*^OL^cKO mice maintained significantly higher excitatory yield across days, indicating a protective or stabilizing effect of increased myelination (p < 0.05, likelihood ratio test with non-overlapping 95% confidence intervals). For putative inhibitory units, WT mice showed higher early yield in cortex than *Fus*^OL^cKO, although these differences did not reach significance in post hoc tests (**Fig. 4C**, p > 0.05, likelihood ratio test with non-overlapping 95% confidence intervals). Within CA1, putative excitatory unit yield was largely stable in both genotypes, with *Fus*^OL^cKO mice exhibiting significantly greater yield than WT across time (**Fig. 4D**, p < 0.05, likelihood ratio test with non-overlapping 95% confidence intervals). Notably, putative inhibitory unit yield in CA1 declined in both groups over time, with WT mice showing higher yield than *Fus*^OL^cKO (**Fig. 4D**, p < 0.05, likelihood ratio test with non-overlapping 95% confidence intervals). Although genotype by day interactions were significant in some comparisons, post hoc tests did not identify consistent differences. These findings indicate that hypermyelination preferentially stabilizes excitatory signals in both cortex and hippocampus, while inhibitory units are less reliably preserved.

To evaluate functional responsiveness at the population level, we computed the absolute signal-to-noise firing rate ratio (|SNFRR|) for putative excitatory and inhibitory units across time and depth. In the cortex, putative excitatory |SNFRR| was stable in *Fus*^OL^cKO mice over the full duration of the study, indicating that hypermyelination did not impact excitatory response amplitude (**Fig. 4E-F**, p > 0.05, likelihood ratio test with non-overlapping 95% confidence intervals). In contrast, WT mice showed significantly greater excitatory |SNFRR| compared to *Fus*^OL^cKO mice (p < 0.05, likelihood ratio test with non-overlapping 95% confidence intervals). For hippocampal CA1, putative excitatory |SNFRR| did not differ between groups and remained stable over time, suggesting region-specific effects of myelination on evoked firing magnitude. Putative inhibitory |SNFRR| in cortex fluctuated greatly within the first few weeks of recording, before decreasing and stabilizing long term (**Fig. 4G-H**). Putative inhibitory |SNFRR| in CA1 declined over time in both groups. *Fus*^OL^cKO mice exhibited higher values early in the recording period, but these differences were not sustained chronically. These data imply that early increases in inhibitory responsiveness associated with hypermyelination are transient and may reflect temporary increases in conduction fidelity or excitability, while long-term benefits are preferentially conferred to excitatory neurons.

### 5.4. Fus^OL^cKO mice demonstrate reduced neuronal firing at the population level in CA1 region of hippocampus

While hypermyelination has been associated with improved conduction velocity and neuronal fidelity, its effects on population-level responsiveness to sensory stimuli remain unclear. To investigate how OL-specific Fus deletion alters evoked ensemble activity, we evaluated multi-unit (MU) firing responses in *Fus*^OL^cKO mice. MU activity reflects the aggregate firing of multiple nearby neurons and provides a measure of population-level response dynamics. We characterized MU responses to visual stimulation using two metrics: MU yield and SNFRR, as previously described (K. Chen et al., 2023b; Moore et al., 2020; Wellman et al., 2020). MU yield was calculated as the percentage of electrode channels exhibiting a significant change in firing rate during a period of drifting grating stimulation compared to an equivalent pre-stimulation period while SNFRR quantified the magnitude of this change for each channel. Both metrics were evaluated across a range of post-stimulus bin sizes (window durations) and latencies (time since stimulus onset) to capture dynamic responses. To visualize temporal response properties, heatmaps of average MU yield were generated across cortical and hippocampal regions (**Fig. 5A**). In both regions, MU responses exhibited a stereotyped onset component within 100 ms post-stimulation, followed by intermittent sustained responses up to 800 ms post-stimulation. This temporal pattern reflects both initial sensory encoding and prolonged circuit reverberations.

**Figure 5.**
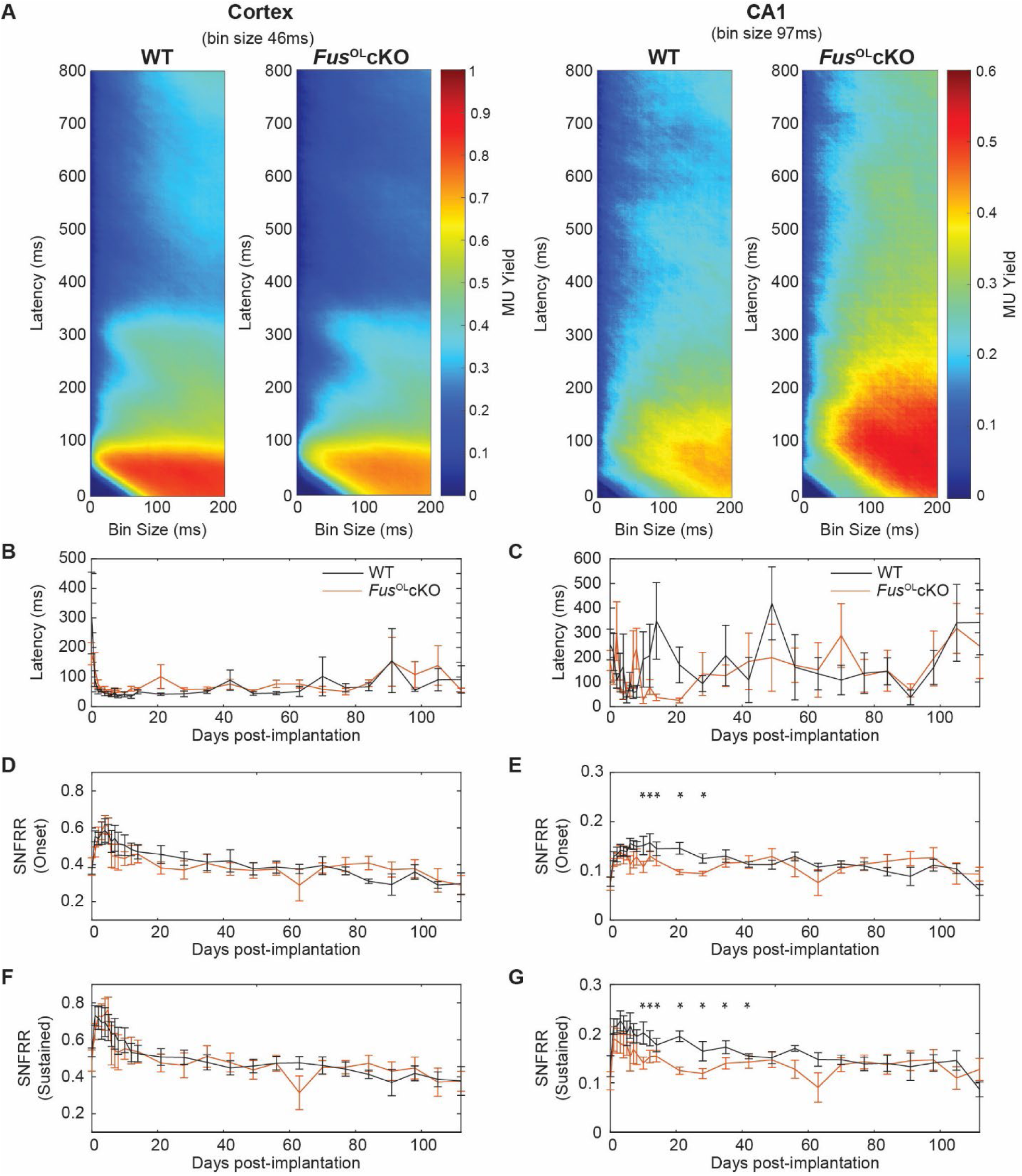
*Fus*^OL^cKO mice demonstrate reduced neuronal firing at the population-level in the CA1 region of the hippocampus. (A) Average of MU yield within the cortex and CA1 region of the hippocampus plotted in 1-ms bins as a function of latency and bin size in response to 1-s of visual stimulation for both *Fus*^OL^cKO mice and WT controls. Visual stimulation evoked a strong, transient multi-unit response within 100ms following stimulation onset, followed by a weaker, sustained MU response between 100 to 800ms. (B) Average latency within the cortex, calculated as the latency at which the optimal MU yield occurred in WT control mice, plotted as a function of time between *Fus*^OL^cKO mice and WT controls. (C) Average latency within the CA1 region of the hippocampus plotted as a function of time between *Fus*^OL^cKO mice and WT controls.<u> No significant differences were observed between groups at either region.</u> (D) Average signal-to-noise firing rate ratio (SNFRR) within the cortex during 100ms onset response to visual stimulation between *Fus*^OL^cKO mice and WT controls. (E) Average SNFRR within the CA1 region of the hippocampus during 100ms onset response to visual stimulation between *Fus*^OL^cKO mice and WT controls. (F) Average SNFRR within the cortex during 100-800ms sustained response to visual stimulation between *Fus*^OL^cKO mice and WT controls. (G) Average SNFRR within the CA1 region of the hippocampus during 100 to 800ms sustained response to visual stimulation between *Fus*^OL^cKO mice and WT controls. **\*** indicates non-overlapping 95% confidence intervals at each time point as determined by likelihood ratio test applied to a linear mixed effects model for WT and *Fus*^OL^cKO mice.

Next, to standardize comparisons, all between-group analyses were anchored at the bin sizes that maximized MU yield in WT mice, corresponding to 46 ms in cortex and 97 ms in CA1, consistent with prior optimization (K. Chen, Cambi, & Kozai, 2023a). This approach ensured sensitivity to stimulus-driven responses while controlling for regional dynamics. To determine whether hypermyelination altered the timing of evoked population responses, latencies were calculated as the time interval between the onset of stimulation and the first statistically measurable change in neuronal firing rate relative to spontaneous activity. We expected that hypermyelination would result in shorter latency durations, indicating accelerated action potential propagation. However, hypermyelination did not significantly affect latency values between WT and *Fus*^OL^cKO mice in the cortex or hippocampus (**Fig. 5B**, p > 0.05, likelihood ratio test with non-overlapping 95% confidence intervals), agreeing with results obtained in previous studies (K. Chen et al., 2023b; Moore et al., 2020; Wellman et al., 2020). These results indicated that the timing of sensory signal arrival and initial encoding was preserved in *Fus*^OL^cKO mice. However, differences emerged in response magnitude. During the onset window (0-100ms post-stimulation), SNFRR was significantly reduced in the CA1 region of *Fus*^OL^cKO mice compared to WT between 10- and 28-d post-implantation (**Fig. 5C**, p < 0.05, likelihood ratio test with non-overlapping 95% confidence intervals) while no differences in onset SNFRR were detected in cortex. During the sustained response window (100–800 ms post-stimulation), CA1 SNFRR remained significantly lower in Fus^OL^cKO mice from 10 to 42 d post-implantation (**Fig. 5D**, p < 0.05, likelihood ratio test with non-overlapping 95% confidence intervals), again with no differences observed in cortex. These findings indicate that although increased myelination in *Fus*^OL^cKO mice does not delay the timing of stimulus-evoked activity, it selectively reduces the magnitude of population responses in hippocampal CA1 without affecting cortical responsiveness. This suggests that hypermyelination may dampen network-level excitability or recurrent activity in hippocampal circuits during sensory processing.

### 5.6. Fus^OL^cKO mice demonstrate reduced laminar connectivity and altered cross-frequency coupling between cortical layers

Myelination affects conduction velocity and synchrony of neural signals across brain regions. Given that previous results showed no significant effects of hypermyelination on single-neuron or population firing properties in cortex, we investigated whether OL-specific *Fus* deletion alters functional connectivity across cortical layers during sensory processing in *Fus*^OL^cKO mice. We used coherence analysis of LFP to quantify the similarity of neural activity between cortical depths. Functional connectivity changes during visual stimulation were expressed as ΔCoherence, comparing evoked versus spontaneous activity across frequency bands and cortical layers. Heatmaps of ΔCoherence revealed prominent differences between *Fus*^OL^cKO mice and WT at 63-d post-implantation between L2/3 and L5 within the delta (2–4 Hz) frequency band (**Fig. 6A**). Over a 16-week period, *Fus*^OL^cKO mice showed significantly reduced normalized coherence between these layers in the delta band from 35- to 77-d post-implantation compared to WT controls (**Fig. 6B**, p < 0.05, likelihood ratio test with non-overlapping 95% confidence intervals). Similarly, significant reductions in coherence were observed between layer 4 and layer 2/3 within the alpha-beta frequency band (8–20 Hz), most pronounced at 5-d post-implantation (**Fig. 6C**, p < 0.05, likelihood ratio test with non-overlapping 95% confidence intervals). This reduction persisted within the first two weeks and again from 28- and 84-d post-implantation in *Fus*^OL^cKO mice compared to WT controls (**Fig. 6D**, p < 0.05, likelihood ratio test with non-overlapping 95% confidence intervals). These findings indicate that hypermyelination in *Fus*^OL^cKO mice significantly impairs the functional communication between key cortical layers across multiple frequency bands.

**Figure 6.**
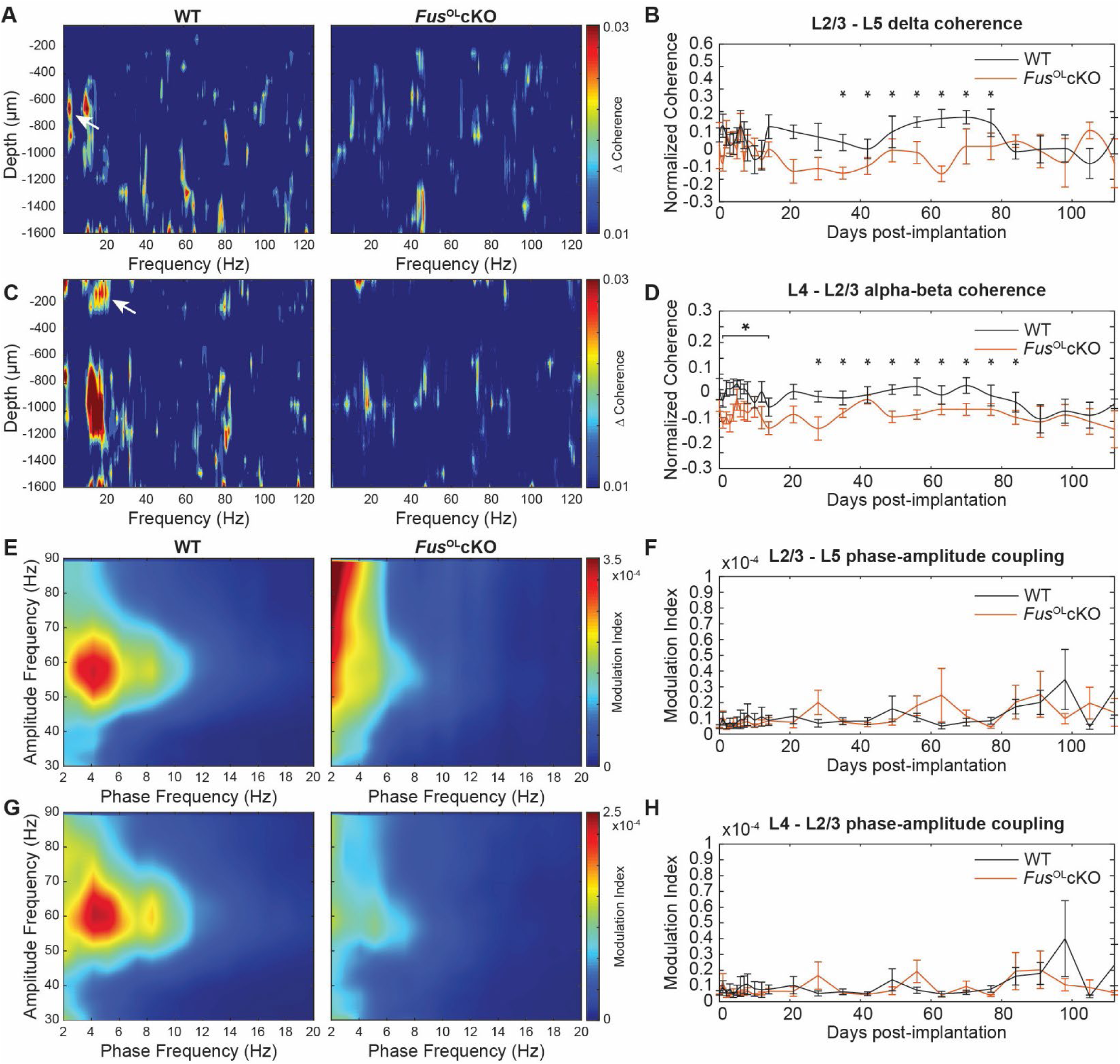
*Fus*^OL^cKO mice demonstrate reduced interlaminar connectivity and altered cross-frequency coupling within the cortex. (A) Heatmap of visually evoked change in coherence (ΔCoherence) as a function of depth and frequency demonstrated large ΔCoherence between supragranular (cortical layer 2/3) and infragranular (cortical layer 5) layers within the delta frequency range between *Fus*^OL^cKO mice and WT controls. (B) Quantification of normalized coherence between supra- and infragranular layers within the delta frequency range over time demonstrates reduced interlaminar coherence in *Fus*^OL^cKO mice compared to WT controls. (C) Heatmap of ΔCoherence as a function of depth and frequency demonstrate large ΔCoherence between supragranular (cortical layer 2/3) and granular (layer 4) within the alpha-beta frequency range between *Fus*^OL^cKO mice and WT controls. (D) Quantification of normalized coherence between granular and supragranular layers within the alpha-beta frequency range over time demonstrated reduced interlaminar coherence in *Fus*^OL^cKO mice compared to WT controls. (E-F) Heatmaps of modulation index (MI) between the phase of low-frequency oscillations and amplitude of high-frequency activity from layer 4 to layer 2/3 in *Fus*^OL^cKO and WT mice. (G) Time course of MI values between layer 4 and layer 2/3 between *Fus*^OL^cKO and WT mice. (H-I) Heatmaps of MI between layer 2/3 and layer 5 between *Fus*^OL^cKO and WT mice. (J) Time course of MI values between layer 2/3 and layer 5 between *Fus*^OL^cKO and WT mice. Both layer 4–2/3 and layer 2/3–5 pathways exhibited reduced theta–gamma coupling in *Fus*^OL^cKO mice, suggesting impaired functional connectivity across cortical layers despite improved single-unit metrics. **\*** indicates non-overlapping 95% confidence intervals at each time point as determined by likelihood ratio test applied to a linear mixed effects model for WT and *Fus*^OL^cKO mice.

Given the critical role of myelin in modulating action potential timing and fidelity, we hypothesized that hypermyelination might influence cross-frequency interactions that facilitate interlaminar communication. We performed phase amplitude coupling (PAC) analysis of LFPs to measure how the phase of slower oscillations modulates the amplitude of faster rhythms across cortical layers, a mechanism implicated in neural coding and network coordination. During visually evoked activity, *Fus*^OL^cKO mice exhibited a lower modulation index (MI) across days for coupling between theta phase (4–8 Hz) and gamma amplitude (55–70 Hz) oscillations between L4 and L2/3, indicating reduced functional connectivity (**Fig. 6E-F**, p < 0.05, likelihood ratio test with non-overlapping 95% confidence intervals). A similar reduction in PAC was found between L2/3 and L5 in the same frequency bands was observed in *Fus*^OL^cKO mice compared to WT (**Fig. 6G-H**, p < 0.05, likelihood ratio test with non-overlapping 95% confidence intervals). Despite the increased SU yield, SNR, and amplitude in the *Fus*^OL^cKO model (**Fig. 1C-F**, **Fig. 2A-C**), this decreased coupling suggests that hypermyelination may enhance synaptic efficiency and reduce redundant or noisy interlaminar signaling. Notably, within the projection from L2/3 to L5 (**Fig. 6G-H**) in *Fus*^OL^cKO mice we saw an increased MI in the lower phase frequency bands (1-4Hz) driving activity within a large range of amplitude frequencies (50–90Hz). This pattern may reflect increased activity of fast-spiking inhibitory interneurons, potentially stabilizing network excitability and preventing hyperexcitation. Together, these results suggest that hypermyelination in *Fus*^OL^cKO mice refines cortical network dynamics by reducing functional connectivity and altering oscillatory interactions between layers, which may improve network efficiency and minimize excitotoxic stress.

### 5.7. Fus^OL^cKO mice retain more neurons during acute injury phase following electrode implantation

To investigate whether hypermyelination influences neuronal survival and circuit integrity following acute brain perturbation, we examined cellular responses in the cortex of *Fus*^OL^cKO and WT mice via immunofluorescence at 2- and 16-wks post-implantation. *Fus*^OL^cKO mice exhibited significantly higher densities of NeuN+ neurons within 100 μm of the electrode track at 2 weeks compared to WT controls (**Fig. 7A**, p < 0.05, one-way ANOVA), indicating enhanced neuronal survival during the acute injury phase. By 16 wks, neuronal densities converged between genotypes, suggesting this protective effect is temporally limited to the early post-implantation period. Despite differences in neuronal survival, axonal density measured by NF200+ fluorescence intensity did not differ significantly between *Fus*^OL^cKO and WT mice at either time point (**Fig. 7B**, p > 0.05, one-way ANOVA). Similarly, counts of CC1+ oligodendrocytes (**Fig. 7C**, p > 0.05, one-way ANOVA) and amount of MBP+ myelin sheaths (**Fig. 7D**, p > 0.05, one-way ANOVA) were comparable between genotypes at both 2 and 16 wks. In order to assess glial immune response, we measured microglial and astrocytic activation near the implanted electrode. Iba-1+ fluorescence intensity, as a marker of microglial activation, did not differ significantly within 150 μm of the electrode between *Fus*^OL^cKO and WT mice at either time point (**Fig. 7E**, p > 0.05, one-way ANOVA). However, GFAP staining revealed a significant reduction in astrogliosis at 2 weeks post-implantation in *Fus*^OL^cKO mice, specifically within 30 μm of the implant site, although this difference was no longer evident at 16 weeks (**Fig. 7F**, p < 0.05, one-way ANOVA). These results suggest that hypermyelination attenuates early astrocytic responses to injury but does not substantially alter microglial activation. Despite using multiple approaches to localize the electrode track in the CA1 hippocampal subregion (900-1600 µm), unequivocal identification was not achieved, precluding direct cellular quantification near the implant in this area. However, immunohistochemical analysis of CA1 pyramidal neurons using MAP2 staining and myelin density with MBP showed no significant genotype differences (data not shown).

**Figure 7.**
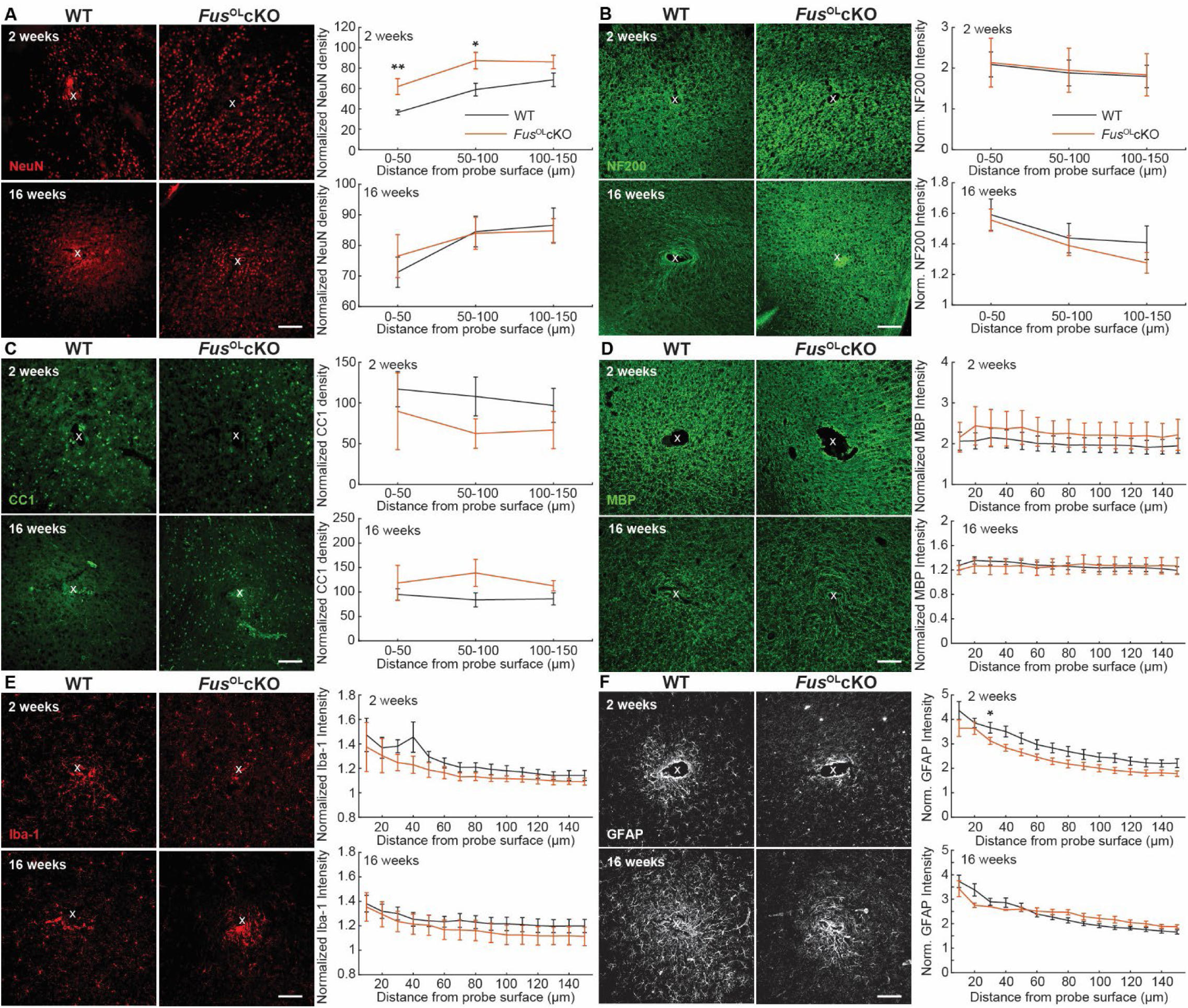
*Fus*^OL^cKO mice demonstrate early retention of neural densities in the cortex within proximity to recording electrode. (A) Representative stain of NeuN+ neurons (*red)* around a microelectrode probe hole within WT and *Fus*^OL^cKO mice at 2-and 16-weeks post-implantation demonstrated acute retention of neuronal densities within 50μm from the probe surface. (B) Representative stain of NF200+ axons (*green*) around a microelectrode probe hole within WT and *Fus*^OL^cKO mice at 2- and 16-weeks post-implantation. (C) Representative stain of CC1+ oligodendrocytes (*green*) around a microelectrode probe hole within WT and *Fus*^OL^cKO mice at 2- and 16-weeks post-implantation. (D) Representative stain of MBP+ myelin (*green*) around a microelectrode probe hole within WT and *Fus*^OL^cKO mice at 2- and 16-weeks post-implantation. (E) Representative stain of Iba-1+ microglia (*red)* around a microelectrode probe hole within WT and *Fus*^OL^cKO mice at 2- and 16-weeks post-implantation. (F) Representative stain of GFAP+ astrocytes (*white*) around a microelectrode probe hole within WT and *Fus*^OL^cKO mice at 2- and 16-weeks post-implantation. Scale bars = 50 μm. * indicates *p*-value < 0.05. ** indicates *p*-value < 0.01.

Glutamatergic neurons constitute the principal excitatory population in CA1 and subiculum, critically shaping hippocampal circuitry (Cembrowski et al., 2018; Ishihara & Fukuda, 2016). To explore mechanisms by which cholesterol-driven hypermyelination might enhance neural function, we quantified glutamatergic synaptic density in CA1 of *Fus*^OL^cKO mice (**Supplementary Fig. 1**). Using postsynaptic density protein 95 (PSD95) as a marker of postsynaptic sites and vesicular glutamate transporter (vGlut1) as a presynaptic excitatory neuron marker, we applied Structured Illumination Microscopy (SIM) and NIS Element software for precise synaptic apposition quantification at 14 days post-implantation (Hong et al., 2017). *Fus*^OL^cKO mice exhibited a significant increase in excitatory synapse number compared to WT controls (p <0.05, one-way ANOVA). These findings link hypermyelination to increased excitatory synaptic connectivity in CA1, which may underlie improved electrophysiological readouts by facilitating stronger excitatory transmission during early post-injury stages.

## 6.0 Discussion

Myelin integrity and OL-mediated metabolic support are fundamental to CA1 circuit function and hippocampal-dependent memory (DeFlitch, Gonzalez-Fernandez, Crawley, & Kang, 2022; Nave & Werner, 2014; Philips & Rothstein, 2017). In *Fus*^OL^cKO mice, conditional deletion of *Fus* in OLs drives cholesterol-mediated hypermyelination (Guzman et al., 2020), allowing us to probe its effects on sensory-evoked activity and network synchrony. Hypermyelination selectively enhanced single-unit detectability and firing rates in CA1 while leaving cortical single-unit responses unchanged, yet it reduced multi-unit population firing, consistent with a shift toward sparser, energy-efficient coding. In the cortex, laminar coherence in delta and alpha–beta bands declined, indicating refined interlaminar synchrony. Post-mortem analysis demonstrated better preservation of NeuN+ neurons in cortex, supporting a neuroprotective role for increased myelination. These results reveal that oligodendrocyte-driven myelin enhancement can optimize individual neuron output and reshape large-scale network coordination without uniformly altering cortical excitability.

### 6.1. Regional specificity of myelin-dependent electrophysiological enhancements

Our results reveal a striking regional specificity in how hypermyelination alters electrophysiological properties. *Fus*^OL^cKO mice exhibit marked improvements in single-unit (SU) detectability and signal-to-noise ratio within hippocampal CA1, effects that persist beyond four weeks of chronic recording (Fig. 2A-E; Fig. 3A-D). By contrast, cortical SU metrics remain unchanged (Fig. 2A-C). These results could be attributed to the heterogeneous nature of myelin distribution in the brain, with myelin density increasing as a function of depth relative to the cortical surface (Tomassy et al., 2014). Potentially, this organization is motivated by hippocampal neurons, especially in CA1, being more reliant on myelin for maintaining excitatory transmission and synaptic plasticity, while cortical neurons may have different compensatory mechanisms or less dependency on myelin for initiating and sustaining their activity (Baltan et al., 2021). Furthermore, improved electrophysiological recordings due to hypermyelination could be due to increased excitability within the hippocampus (Pinatel et al., 2023). *Fus*^OL^cKO mice displayed a significantly higher density of excitatory synapses within hippocampal CA1 (Supplementary Fig. 1), suggesting that hypermyelination strengthens synaptic connectivity.

Paradoxically, multi-unit (MU) firing rates and signal-to-noise firing rate ratios in CA1 are reduced during both onset (0–100 ms) and sustained (100–800 ms) responses (Fig. 5C–D). This shift toward sparser MU activity, despite elevated SU performance, is consistent with an energy-efficient coding strategy supported by myelin’s reduction of ionic flux and metabolic cost (Attwell & Laughlin, 2001; Suminaite, Lyons, & Livesey, 2019). This paradoxical reduction in population-level firing may reflect enhanced energy efficiency. By facilitating faster, more reliable axonal conduction, myelin allows individual neurons to maintain output with fewer synchronous spikes, thereby reducing overall metabolic demand during network activity and increasing network efficiency (Gerevich et al., 2023). Interestingly, the region-specific efficacy of *Fus*^OL^cKO hypermyelination in CA1 but not cortex contrasts with prior findings using clemastine, which improved electrophysiological signals in both regions (K. Chen et al., 2023b). This divergence suggests that cortical resilience may depend more on non-myelinating functions of oligodendrocytes, such as metabolic support or neuroimmune modulation, whereas CA1 appears more sensitive to alterations in myelin thickness itself.

### 6.2. Hypermyelination preserves chronic single-unit fidelity and neuronal survival in CA1

Hypermyelination in *Fus*^OL^cKO mice supports long-term neuronal preservation and recording stability following chronic microelectrode implantation in hippocampal CA1. Post-mortem analysis showed increased NeuN+ neuronal density and preserved NF200+ axonal integrity (Fig. 7A-B), indicating that hypermyelination may confer neuroprotection against implantation-induced injury, which is known to degrade neuronal viability and activity (T. D. Kozai, Gugel, et al., 2014; T. D. Kozai, Li, et al., 2014; Wellman et al., 2019). This protection likely sustains single-unit fidelity through improved axonal conduction and structural support for neuronal processes. Preservation of excitatory synapse density (Supplementary Fig. 1) further supports synaptic integrity as a substrate for stable network function.

Recording performance in the hippocampus typically suffers due to electrode penetration through heavily myelinated axon tracts projecting to CA1 pyramidal neurons. Preliminary data suggest that greater myelin presence helps maintain axonal functionality and synaptic connectivity, which is notable given the known difficulty in maintaining chronic CA1 recordings beyond one month (T. D. Kozai et al., 2015). This limitation restricts studies of memory, learning, and plasticity in a region critical for cognitive function. *Fus*^OL^cKO mice show significantly improved recording performance in the hippocampus (Fig. 5), highlighting myelin’s role in mitigating chronic recording failure. These observations align with the concept that myelin enhances axonal efficiency by reducing sodium influx during action potentials and lowering ATP demands for repolarization (Harris & Attwell, 2012).

*Fus*^OL^cKO hypermyelination selectively increases the percent of myelinated small diameter-axons (Guzman et al., 2020), a population characterized by high membrane surface to- volume-ratios and correspondingly greater metabolic demand during rapid firing (Harris & Attwell, 2012; Suminaite et al., 2019). These fine-caliber fibers are essential for precise spike timing and local circuit computation in CA1,yet are especially vulnerable to energy depletion in zones of sparse vascularization and elevated synaptic throughput (Ji et al., 2021; Zhang et al., 2019). Microelectrodes are typically implanted from the dorsal surface, traversing axon-rich regions such as the stratum oriens and alveus, which contain myelinated Schaffer collateral projections from CA3 and extrahippocampal inputs; this trajectory disrupts and demyelinates axons critical for synaptic drive into CA1. By adding extra insulation to these metabolically sensitive axons, hypermyelination in *Fus*^OL^cKO mice reduces ionic flux and ATP requirements per action potential (Harris & Attwell, 2012). In parallel, oligodendrocyte-mediated transfer of glycolytic lactate and glucose via MCT1 and GLUT1 transporters provides an auxiliary substrate to buffer metabolic stress (Funfschilling et al., 2012; Saab et al., 2016). This dual enhancement of structural insulation and glial metabolic support likely underpins the observed preservation of neuronal survival and single-unit fidelity under conditions of chronic implant stress.

Despite reduced population-level firing in CA1, the preservation of excitatory unit yield indicates more stable and reliable signal detection relative to wild-type controls. Hypermyelination correlates with increased neuronal survival during the acute injury phase near the implant, contributing to sustained recording quality. Functional connectivity analyses reveal reduced laminar coherence and altered cross-frequency coupling in cortical layers, which may signify improved network efficiency (Figs. 2-5). These results demonstrate that hypermyelination in *Fus*^OL^cKO mice counters typical chronic recording degradation in CA1, a region vital for hippocampal-dependent cognitive processes. These findings extend prior pharmacological studies with clemastine (K. Chen et al., 2023b) by isolating the contribution of myelin thickness itself from broader oligodendrocyte effects (Guzman et al., 2020), emphasizing the importance of myelin integrity for sustaining hippocampal circuit function under chronic stress.

### 6.3. Refinement of cortical network synchrony by hypermyelination

In *Fus*^OL^cKO mice, cortical single-unit and multi-unit activity metrics remained stable (Figs. 2, 3), yet interlaminar coherence was significantly reduced in delta (2–4 Hz) and alpha–beta (8–20 Hz) frequency bands (Fig. 6A-D). Additionally, phase–amplitude coupling between theta and gamma oscillations across L4-L2/3 and L2/3-L5 was diminished (Fig. 6E-J). These findings indicate that hypermyelination modulates the temporal coordination of oscillatory activity between cortical layers without broadly enhancing cortical excitability. Global oscillatory power across frequency bands was unchanged, consistent with prior observations that oligodendrocyte loss or pharmacological rescue primarily affect the strength of specific neuronal oscillations (K. Chen et al., 2023b; Wellman et al., 2020). Our results suggest that hypermyelination modulates the activity of individual neurons without proportionally increasing synchrony across large neuronal populations. Hypermyelination may fine-tune local circuit dynamics while preserving the overall balance of cortical oscillations (Pajevic, Basser, & Fields, 2014). Notably, *Fus*^OL^cKO mice showed reduced laminar connectivity relative to wild-type controls, suggesting that increased myelin thickness strengthens single-neuron function but may impair coordinated communication between cortical layers. Rather than facilitating intra- and interlaminar signaling, hypermyelination could cause temporal misalignment across cortical circuits, potentially disrupting the integration of complex information (Pajevic et al., 2014). This nuanced modulation contrasts with the preservation of firing properties at the single-neuron level, implying that hypermyelination selectively influences the coordination of population activity across cortical layers. Such changes in cortical network dynamics may impact cognitive functions that depend on precise interlaminar signaling, including sensory integration and higher-order processing (Chadwick et al., 2023; Wimmer et al., 2015).

## 7.0 Conclusion

This study advances understanding of how hypermyelination preserves hippocampal circuit function by supporting conduction velocity, and synaptic integrity within CA1. These effects contribute to stable single-unit activity and sustained network function, which are essential for memory encoding and cognitive processing. Such preservation is particularly relevant in aging and neurodegenerative diseases where disrupted myelin impairs cognitive functions (J. F. Chen et al., 2021; Su et al., 2020). Oligodendrocyte-mediated myelin remodeling emerges as a critical modulator of circuit resilience and plasticity, influencing excitatory–inhibitory balance and protecting neuronal populations against chronic injury (J. F. Chen et al., 2021). These glial functions likely extend beyond classical insulation roles to include metabolic and trophic support essential for maintaining long-term neural network stability (Wellman et al., 2018). From a translational neuroscience perspective, targeting myelin health may offer therapeutic avenues to restore or enhance cognitive functions affected by demyelinating or neurodegenerative pathology (Takashi D. Y. Kozai, Jaquins-Gerstl, Vazquez, Michael, & Cui, 2015; Salatino, Ludwig, Kozai, & Purcell, 2017; Schwartz, Cui, Weber, & Moran, 2006). Moreover, maintaining myelin integrity could improve the fidelity of electrophysiological signals used to study circuit dynamics in vivo, facilitating more accurate investigation of learning and memory mechanisms (Wellman et al., 2018; Wellman & Kozai, 2017). Future research should address how modulating myelin plasticity influences synaptic modifications and network oscillations in disease and recovery contexts.

## Acknowledgement

We thank Anthony Vetter software specialist at Nikon for his help in developing a workflow in the General Analysis (GA3) software to quantify the appositions of pre- and postsynaptic puncta. This work was supported by: NIH R21NS108098, NIH R01NS094396, NIH R01NS105691, NIH R01NS115707, NIH R03AG072218, NIH R01NS129632, F99NS124186, and NSF CBET CAREER 1943906.

---

**Supplementary Figure 1.**
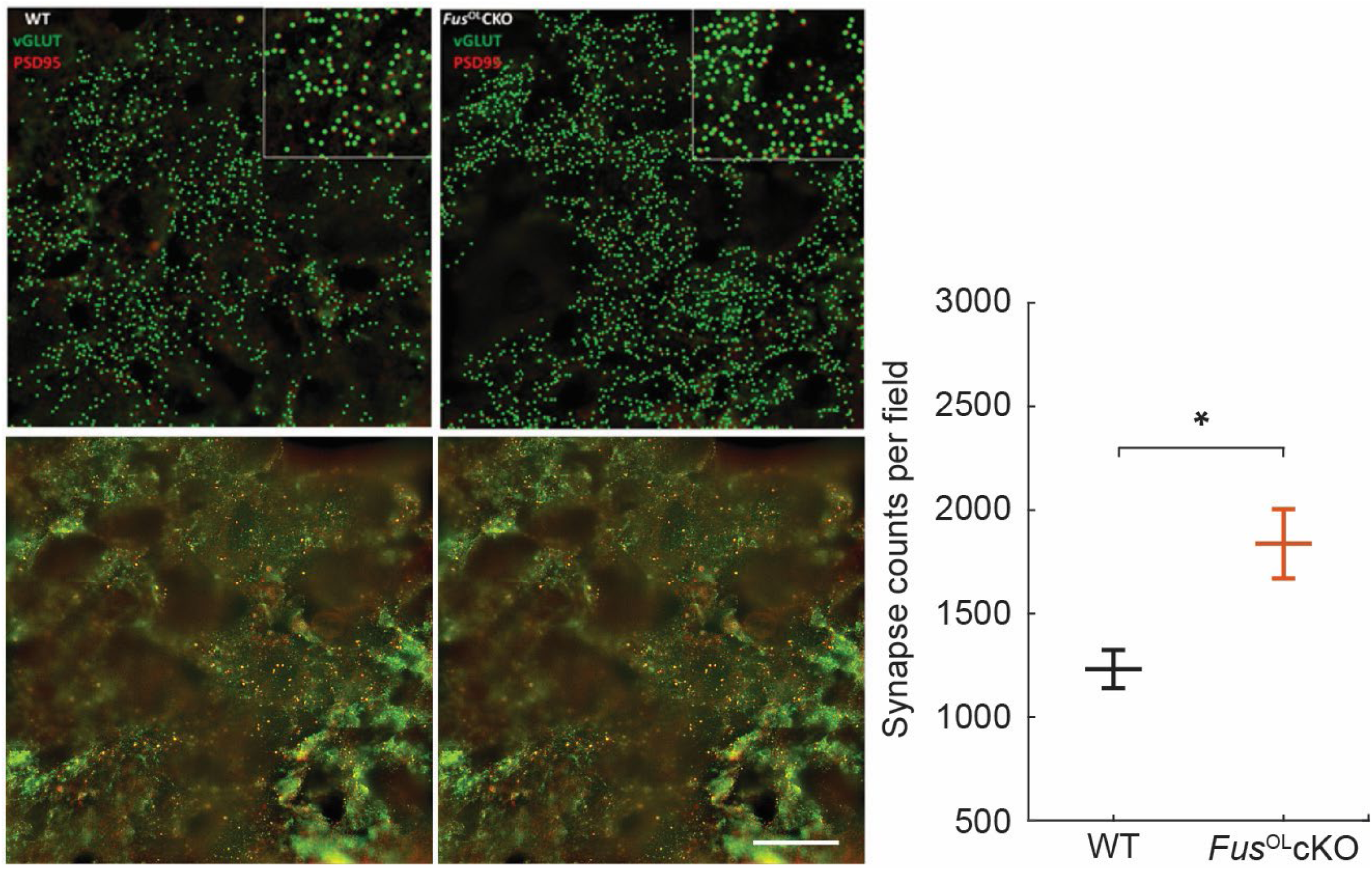
*Fus*^OL^cKO mice exhibit a greater number of excitatory synapses in CA1 subregion compared to WT controls. *Top:* xNIS Element rendition of postsynaptic density protein 95 (PSD95, *red*) puncta, a marker of excitatory postsynaptic sites, and of vesicular glutamate transporter (vGlut1, *green*) puncta, a presynaptic excitatory neuron marker. *Bottom*: Representative confocal images of horizontal tissue sections co-stained with PSD95 and vGlut1 taken at 100X magnification to compare the number of excitatory synapses within CA1 region of hippocampus between *Fus*^OL^cKO mice and WT controls. Graph shows synapse counts/100x oil Structured Illumination Microscopy reconstructed field based on the number of pre- and post-synaptic appositions per field of view. Scale bar = 10 μm. * indicates *p*-value < 0.05.

