## SupplementalFigures for "Hypermyelination Improves Strength and Detection of Neuronal Activity in the CA1 Hippocampus and Facilitates Neuroprotection in *Fus*^OL^cKO Mice"

**Abbreviated title:** Myelin Enhances CA1 Neuronal Recordings in *Fus*$^{OL}$cKO Mice

**Authors:** Steven M. Wellman[1,2], Kelly Guzman[3,4], Naofumi Suematsu[1,2,8], Teresa Thai[1,2], Te-Hsuan Tung[4], Camila Garcia Padilla[1,2], Sadhana Sridhar[7], Keying Chen[1,2], Franca Cambi[3,4], Takashi D.Y. Kozai [1,2, 5, 6,7]*

1) Department of Bioengineering, University of Pittsburgh, Pittsburgh, PA, USA, 15213
2) Center for Neural Basis of Cognition, Pittsburgh, PA, USA, 15213
3) Veterans Administration Pittsburgh, Pittsburgh, PA, USA, 15213
4) Department of Neurology, University of Pittsburgh, Pittsburgh, PA, USA, 15213
5) Center for Neuroscience, University of Pittsburgh, Pittsburgh, PA, USA, 15213
6) McGowan Institute of Regenerative Medicine, University of Pittsburgh, Pittsburgh, PA, USA, 15213
7) NeuroTech Center, University of Pittsburgh Brain Institute, Pittsburgh, PA, USA, 15213
8) Department of Molecular Cell Physiology, Kyoto Prefectural University of Medicine, Kyoto, Japan, 602-8566

**Number of pages:** 25
**Number of figures:** 7 main figures, 1 supplemental figure

**Acknowledgement:** We thank Anthony Vetter software specialist at Nikon for his help in developing a workflow in the General Analysis (GA3) software to quantify the appositions of pre- and postsynaptic puncta. This work was supported by: NIH R21NS108098, NIH R01NS094396, NIH R01NS105691, NIH R01NS115707, NIH R03AG072218, NIH R01NS129632, F99NS124186, and NSF CBET CAREER 1943906.

**Supplemental Material**

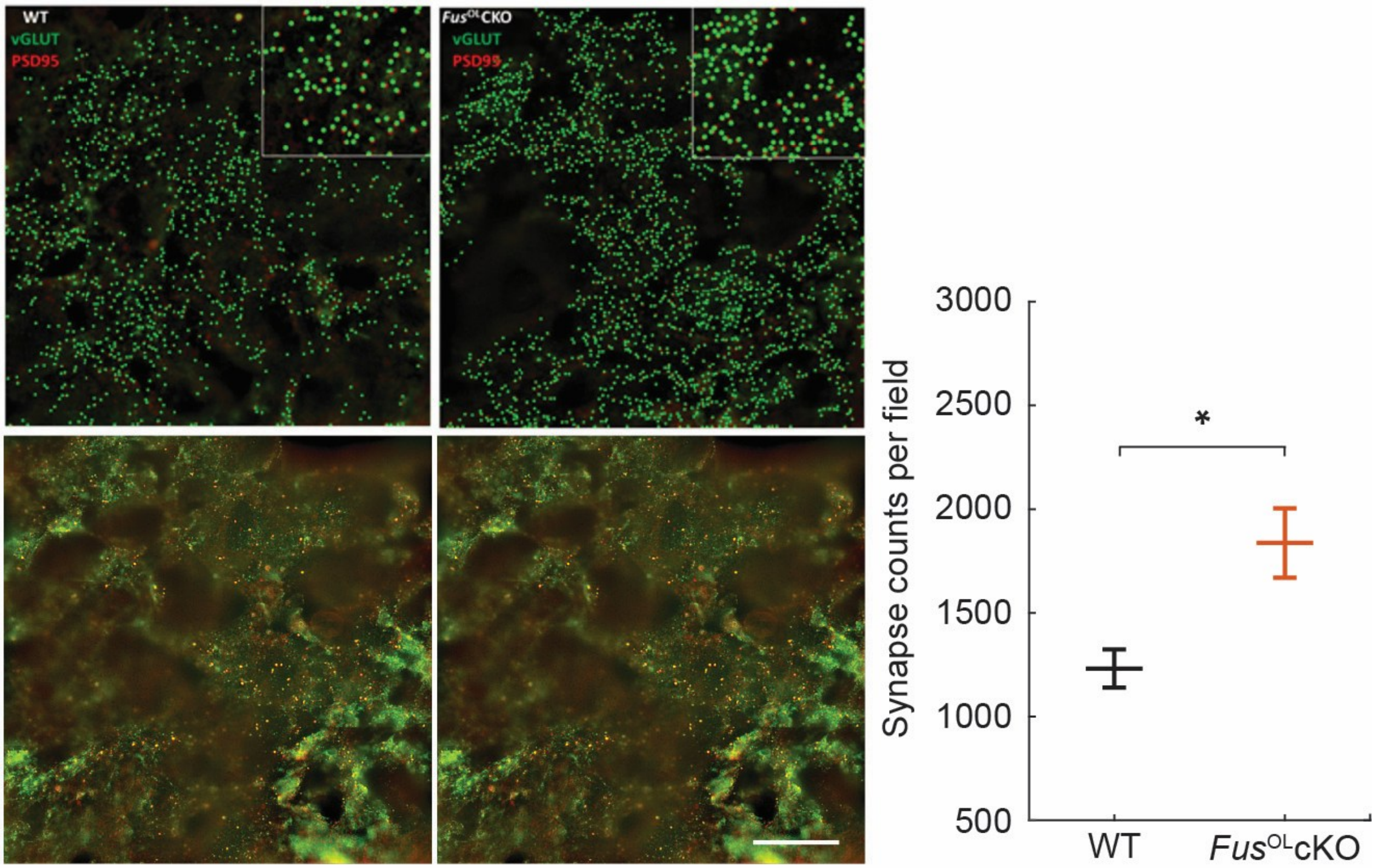


**Supplementary Figure 1. *Fus*$^{OL}$cKO mice exhibit a greater number of excitatory synapses in CA1 subregion compared to WT controls.** *Top:* xNIS Element rendition of postsynaptic density protein 95 (PSD95, *red*) puncta, a marker of excitatory postsynaptic sites, and of vesicular glutamate transporter (vGlut1, *green*) puncta, a presynaptic excitatory neuron marker. *Bottom*: Representative confocal images of horizontal tissue sections co-stained with PSD95 and vGlut1 taken at 100X magnification to compare the number of excitatory synapses within CA1 region of hippocampus between *Fus*$^{OL}$cKO mice and WT controls. Graph shows synapse counts/100x oil Structured Illumination Microscopy reconstructed field based on the number of pre- and post-synaptic appositions per field of view. Scale bar = 10 μm. * indicates $p$-value < 0.05.

### REFERENCES
